# Testicular macrophages are recruited during a narrow time window by fetal Sertoli cells to promote organ-specific developmental functions

**DOI:** 10.1101/2022.05.05.490754

**Authors:** Xiaowei Gu, Anna Heinrich, Tony DeFalco

## Abstract

While macrophages are most commonly known for their roles in innate immunity, a growing body of evidence supports the idea that fetal-derived tissue-resident macrophages play developmental roles during organogenesis. In the testis, it has long been proposed that macrophages are important players in steroidogenesis and other testicular functions, but which macrophage populations are involved is unclear. We previously showed that macrophages play critical roles in fetal testis morphogenesis and reported the presence of 2 unique adult testicular macrophage populations, interstitial and peritubular. There has been some debate regarding the hematopoietic origins of testicular macrophages and whether distinct macrophage populations promote specific testicular functions. Here we have undertaken an extensive lineage-tracing study of mouse hematopoietic cells. We found that, while yolk-sac-derived macrophages comprise the earliest testicular macrophages, fetal hematopoietic stem cells (HSCs) give rise to monocytes that colonize the gonad during a narrow time window in mid-gestation, after which time HSCs no longer contribute to testicular macrophages. These long-lived monocytes, over the course of fetal and postnatal life, differentiate into testicular macrophages. Our data indicate that Sertoli cells, and not germ cells, are required for recruitment of immune cells and peritubular macrophage differentiation. Finally, we show that yolk-sac-derived macrophages and HSC-derived macrophages play distinct roles in testis cord morphogenesis, whereas interstitial macrophages promote adult Leydig cell proliferation and steroid production. Overall, our findings offer clarity regarding the origins of testicular macrophages and provide insight into the diversity of their tissue-specific developmental roles.

## Introduction

Macrophages are ubiquitous throughout the body and play tissue-specific roles, often with unique functions critical for organ development and homeostasis (1, 2). While traditionally appreciated for their importance in the innate immune response, macrophages also have unexpected functions that are dramatically distinct from their classic phagocytic and antigen-presenting roles. Most aspects of macrophage biology that have been studied in the literature are either in postnatal or adult contexts, but recent research in the field has started to focus on the hematopoietic origins of macrophages during embryogenesis and their potentially diverse repertoire of cellular functions during fetal development (3, 4). A major open question is whether hematopoietic origins dictate the capability of macrophages to perform tissue-specific activities, or if the cellular microenvironment in which macrophages reside is the major driving factor underlying tissue-resident macrophage phenotypes and activities.

Using a number of specific labeling and targeting tools in the mouse model system, the hematopoietic origin of macrophages within mammalian tissues has recently been an active area of research (3, 4). Numerous reports have shed light onto new models of hematopoiesis that diverge from a classical model in which all macrophages were assumed to differentiate from circulating monocyte progenitors originating from adult bone marrow hematopoietic stem cells (HSCs) in what is termed definitive hematopoiesis (5). Studies over the past 15 years have revealed that, in contrast to the previous dogma centered around adult definitive hematopoiesis, embryonic hematopoiesis is a major contributor to tissue-resident macrophages within adult organs (6–12). Furthermore, it has also been shown that a number of organs, such as the brain, maintain their tissue-resident macrophage populations (microglia) independently of bone-marrow-derived HSCs, while other organs, such as the gut, exhibit a heavy reliance on bone-marrow HSC-derived monocytes to maintain homeostasis of their tissue-resident macrophages (4). It is clear that each organ has unique dynamics and kinetics of macrophage colonization and homeostasis during development; however, the underlying mechanisms that drive these organ-specific phenotypes are unclear.

Mammalian hematopoiesis occurs in several waves to give rise to cells that seed organs and differentiate into tissue-resident macrophages (3,4,13). The first wave, called primitive hematopoiesis, takes place in the embryonic yolk sac (YS) blood islands starting at E7.0 in mice, and gives rise to primitive YS-derived macrophages. Between E8.0 and E8.5, a second wave of hematopoiesis from YS hemogenic endothelium, sometimes referred to as the transient definitive wave, gives rise to a class of progenitors called erythromyeloid precursors (EMPs). After blood circulation ensues starting at E8.5, EMPs migrate to the fetal liver and expand in number to differentiate into monocytes and other hematopoietic lineages. Soon thereafter, within the hemogenic endothelium of the embryo proper, immature HSCs arise in the para-aortic splanchnopleura (P-Sp) region and give rise to fetal HSCs at E10.5 within the aorta-gonad-mesonephros region (AGM). These HSCs then colonize the fetal liver to initiate steady-state definitive hematopoiesis in the embryo; starting at E12.5, the fetal liver is the main site of hematopoiesis where myeloid and lymphoid cells of the nascent embryonic immune system will be generated. By the end of embryogenesis, fetal liver HSCs will seed the fetal bone marrow, where these fetal HSCs will give rise to adult bone marrow HSCs.

The gonad, and the testis in particular, is a unique immune environment where inflammatory responses are generally suppressed to protect gametes from attack by the immune system and to ensure continued fertility (14–16). A major component of this specialized immune environment is the testicular macrophage, which is a central contributor to testicular immune function (17–19) and comprises a significant proportion (up to 25%) of the interstitial compartment of the testis (20). We have previously shown that macrophages are present in the early undifferentiated fetal gonad and YS hematopoietic precursors give rise to macrophages in the nascent fetal testis (21), and other recent reports have also examined the origins of testicular immune cells, implicating monocytes and other myeloid cell types (22–24). In terms of function, we demonstrated that fetal testicular macrophages are required for testis morphogenesis during initial gonadogenesis, likely through modulation of vascular development that is critical for the formation of testicular architecture (21, 25).

By adulthood, at least 2 major testicular macrophage populations have been described, based on cell surface marker expression and localization within the organ: CSF1R+CD206+MHCII- interstitial macrophages, which are embedded amongst Leydig cells and blood vessels in the interstitial parenchyma; and CSF1R- CD206-MHCII+ peritubular macrophages, which are exclusively localized to the surface of seminiferous tubules (23, 26). There has been some debate regarding the hematopoietic origin of adult testicular macrophage populations, with one study proposing a circulating, bone-marrow-HSC-derived origin for peritubular macrophages, and others proposing a fetal-monocyte-derived origin for these cells (22–24). Testicular macrophages have long been implicated in regulating steroidogenesis by testicular Leydig cells (27–29), while peritubular macrophages have been proposed to promote spermatogonial differentiation (26); however, specific functions for distinct testicular macrophage populations have not been functionally investigated.

Here we have investigated in detail the hematopoietic origins and functions of testicular macrophages. Using a wide array of lineage-tracing tools in mice, we found that, while macrophages of the early nascent testis are YS-derived, monocytes originating from AGM-derived HSCs recruited during a specific embryonic time window are the major hematopoietic precursor to both adult testicular macrophage populations. We also confirm that YS-derived macrophages are a minor contributor to adult testicular macrophages. Our analyses revealed that Sertoli cells are required to recruit monocytes into the fetal testis and germ cells are dispensable for differentiation and localization of adult interstitial and peritubular macrophages. We specifically ablated YS macrophages and found that testicular cord morphogenesis was disrupted and ectopic monocytes migrated into the testis; our data suggest that distorted myeloid cell ratios specifically impact testis cord morphogenesis but do not disrupt Sertoli cell gene expression. Finally, we used *in vitro* assays to demonstrate that testicular interstitial macrophages promote Leydig cell proliferation and steroidogenesis, providing new, specific insights into previous long-standing observations in the field. Our findings highlight the complexity of testicular macrophage origins and functions, and may provide new perspective into mechanisms that underlie gonadal developmental defects and infertility.

## Results

### Csf1r+ fetal definitive progenitors give rise to adult testicular macrophages

Given that *Csf1r*-expressing cells in the yolk sac (YS) appear at embryonic day 7.5 (E7.5) (8), representing the earliest primitive macrophage progenitors, and peak between E9.5 and E10.5 (30), induction with 4- hydroxytamoxifen (4OHT) at E8.5 in the *Csf1r-*creER model can specifically label YS-derived EMPs (31, 32). The fetal liver from E10.5 to E12.5 then becomes the main site of *Csf1r*-expressing progenitors that have erythroid and myeloid potential (33), emerging later than YS EMPs. To identify whether YS-derived EMPs or subsequent *Csf1r*+ definitive progenitors contribute to fetal and postnatal testicular macrophages, we crossed *Csf1r-*creER mice with *Rosa-*Tomato reporter mice and delivered a single injection of 4OHT at E8.5, E10.5 or E12.5 to induce the activity of *Csf1r-*creER, resulting in irreversible Tomato expression in *Csf1r*-expressing cells and their progeny (Fig. 1A).

**Figure 1.**
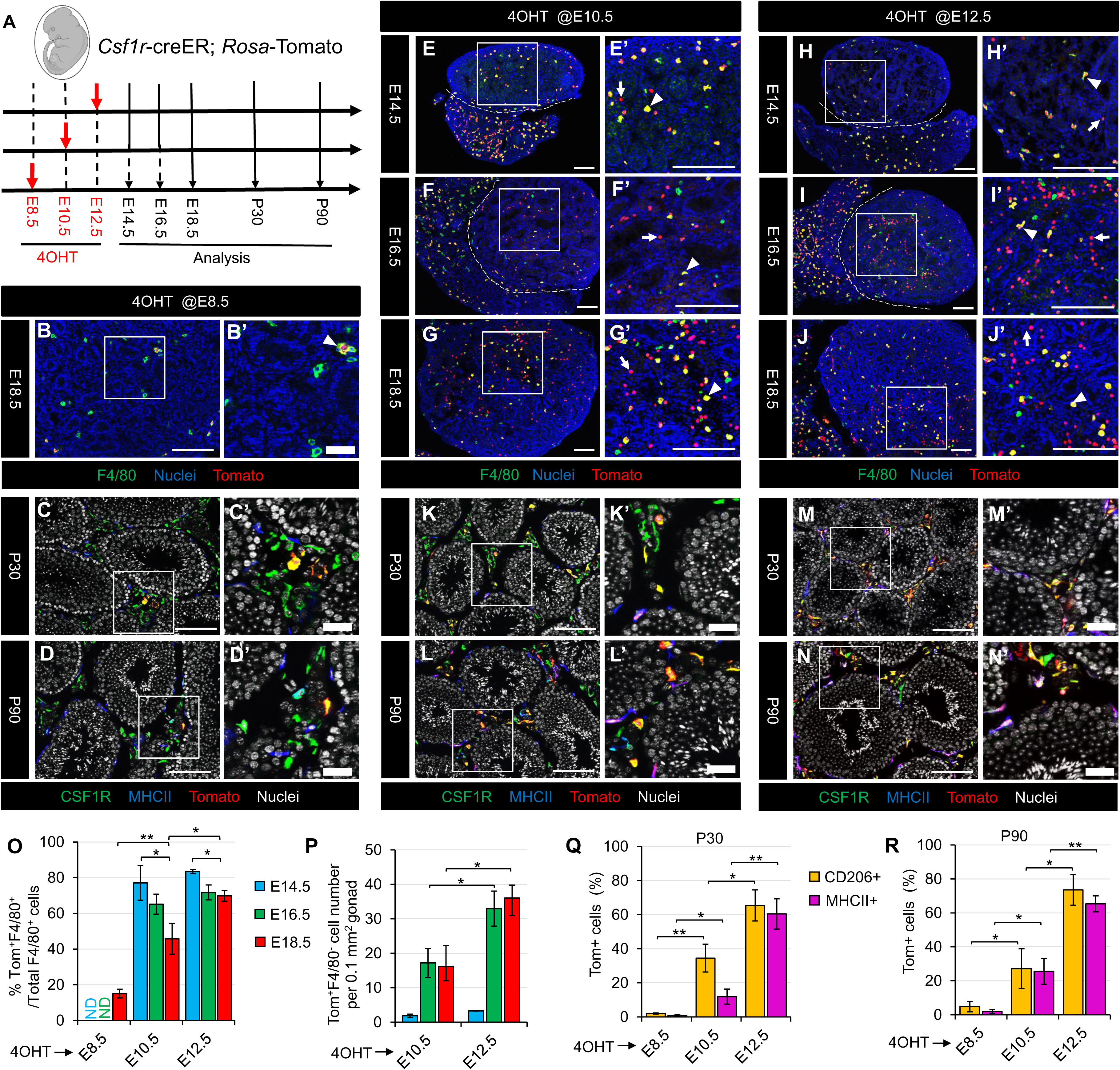
*Csf1r*-expressing fetal definitive progenitors give rise to adult testicular macrophages. (A) creER; *Rosa*-Tomato embryos and juvenile/adult mice. (B-N) Images of testes at various stages from *Csf1r*- creER; *Rosa*-Tomato mice exposed to 4OHT at E8.5 (B-D), E10.5 (E-G,K,L), or E12.5 (H-J,M,N). In all figures throughout this study, prime figures (e.g., A’ relative to A) are higher-magnification images of the boxed regions in the image to their left. Dashed lines indicate gonad-mesonephros boundary. Arrowheads denote Tomato-expressing F4/80+ macrophages and arrows denote Tomato-expressing F4/80-negative cells. Thin scale bar, 100 μm; thick scale bar, 25 μm. (O-N) Graphs showing quantification (*n*=3) of percent Tomato-expressing F4/80+ macrophages at E14.5, E16.5, or E18.5 (O), number of Tomato-expressing F4/80- cells per unit area at E14.5, E16.5, or E18.5 (P), and percent Tomato-expressing interstitial (CD206+) and peritubular (MHCII+) macrophages at P30 (Q) or P90 (R) in *Csf1r*-creER; *Rosa*-Tomato testes induced with 4OHT at various embryonic stages. Data are shown as mean +/- SD. \**P*<0.05; \*\**P*<0.01 (two-tailed Student’s *t*-test).

When pulsed with 4OHT at E8.5, as a positive control, we examined microglia in E18.5 and postnatal brains, which are derived exclusively from YS progenitors (7,11,31). Extensive Tomato+ cells colocalized with F4/80 throughout the brain as expected (Fig. S1A, S1B and S1M), whereas we detected only a few Tomato- labeled F4/80+ testicular macrophages at E18.5 (Fig. 1B and 1O), even fewer CSF1R^+^MHCII^-^ or CD206^+^MHCII^-^ interstitial macrophages and CSF1R^-^MHCII^+^ or CD206^-^MHCII^+^ peritubular macrophages with Tomato at postnatal day 30 (P30) and P90 (Fig. 1C, 1D, 1Q and 1R), suggesting that YS-derived EMPs had a minor contribution to testicular macrophages. We also could induce labeling of microglia as efficiently at E10.5 (>80%) as at E8.5, with no dilution from fetal to adult stages, but E12.5 induction only labeled about 40% of microglia (Fig. S1C-F, S1M), confirming an earlier report that YS macrophage progenitors travel to tissues within a restricted time window (30). However, while Tomato-labeled liver macrophages (Kupffer cells) were diluted from fetal to adult stages when induced at E8.5 or E10.5 (Fig. S1G-J, S1N), a large number of Kupffer cells were stably labeled when induced at E12.5 (Fig. S1K, S1L and S1N). These findings indicate that EMP-derived Kupffer cells are partially replaced by *Csf1r*^+^ definitive progenitor-derived macrophages, supporting a previous study proposing Kupffer cells come from two sources: YS-derived EMPs and fetal immature HSCs (7, 34). These data suggest that *Csf1r*^+^ macrophage progenitors produced at E12.5 are no longer largely derived from the YS. Similar to Kupffer cells, when pulsed at E10.5 and E12.5, the majority of F4/80^+^ macrophages in E14.5, E16.5, and E18.5 testes expressed Tomato (Fig. 1E-J and 1O). Although there was a downward trend in the percentage of Tomato+ testicular macrophages in E18.5 testes compared with E14.5 testes (Fig. 1O), there was still a significant increase as compared to induction at E8.5, especially induction at E12.5 (Fig. 1O). Additionally, high numbers of Tomato-labeled interstitial and peritubular macrophages in P30 and P90 testes were detected when induced at E10.5 or E12.5, and their frequency had the most substantial increase in E12.5-induced P30 and P90 testes (Fig. 1K-N, 1Q and 1R).

Of note, we found that induction at E10.5 yielded some Tomato^+^F4/80^-^ cells in testes starting from E16.5 (Fig. 1E-G and 1P), whereby induction at E12.5 had more such cells emerging in late fetal testes and liver (Fig. 1H-J and 1P). When induced at E10.5 or E12.5, all Tomato+ cells in E18.5 liver and testes were positive for CD45 (a pan-leukocyte marker) (Fig. S2A-D), but only a few KIT (a HSC marker)-positive cells in the liver expressed low levels of Tomato (Fig. S2E and S2F), suggesting *Csf1r-*creER-driven Tomato-labeled progenitors may be the progeny of fetal HSCs. In addition, some GR1^+^ cells, primarily representing granulocytes and monocytes, were also labeled in E18.5 testes and liver (Fig. S2G-J). Given lymphoid cells, particularly T cell subsets, emerging and gradually increasing in postnatal testes with age (35), these data imply that the CD45^+^Tomato^+^F4/80^-^ cell population in later fetal testes belongs to the myeloid lineage. To further verify whether *Csf1r*^+^ progenitors have lymphoid potential, we examined Tomato expression in adult spleen. We found that CD4^+^ T cells and B220^+^ B cells were labeled by Tomato in E10.5 or E12.5-induced adult spleen, and E12.5 induction resulted in extensive labeling (Fig. S3A-F), suggesting that *Csf1r-*creER-driven Tomato expression was activated in lymphoid cells when induced at E10.5 and E12.5. However, almost no GR1^+^ cells were labeled in adult spleen (Fig. S3G-I); this lack of labeling may be related to the short circulatory lifespan (less than 1 day in mice and 5.4 days in humans) and fast turnover rate of mature neutrophils (36). Together, these results show that *Csf1r*^+^ definitive progenitors that contribute to testicular macrophage pools are likely derived from early fetal HSCs rather than YS-derived EMPs.

### The contribution of AGM-derived HSCs to adult testis macrophages has a unique recruitment window

Although the *Csf1r*-inducible fate mapping model likely targets fetal HSC-derived multipotent progenitors that give rise to tissue-resident macrophages according to the above-described results, *Csf1r* is also expressed in mature macrophages (37, 38), complicating interpretation of the results. Given that KIT (also called C-KIT or CD117) is widely expressed in fetal and adult HSCs as well as in early YS and fetal liver progenitors, but not in mature hematopoietic cells (39), we used the *Kit*^creER^ mouse strain to trace hematopoietic output from hematopoietic progenitors and HSCs. To test whether fetal HSCs contribute to testicular macrophages, we administered single 4OHT injections at different time points during gestation (E8.5, E10.5, E12.5, E14.5) to label *Kit*^creER^; *Rosa-*Tomato embryos (Fig. 2A).

**Figure 2.**
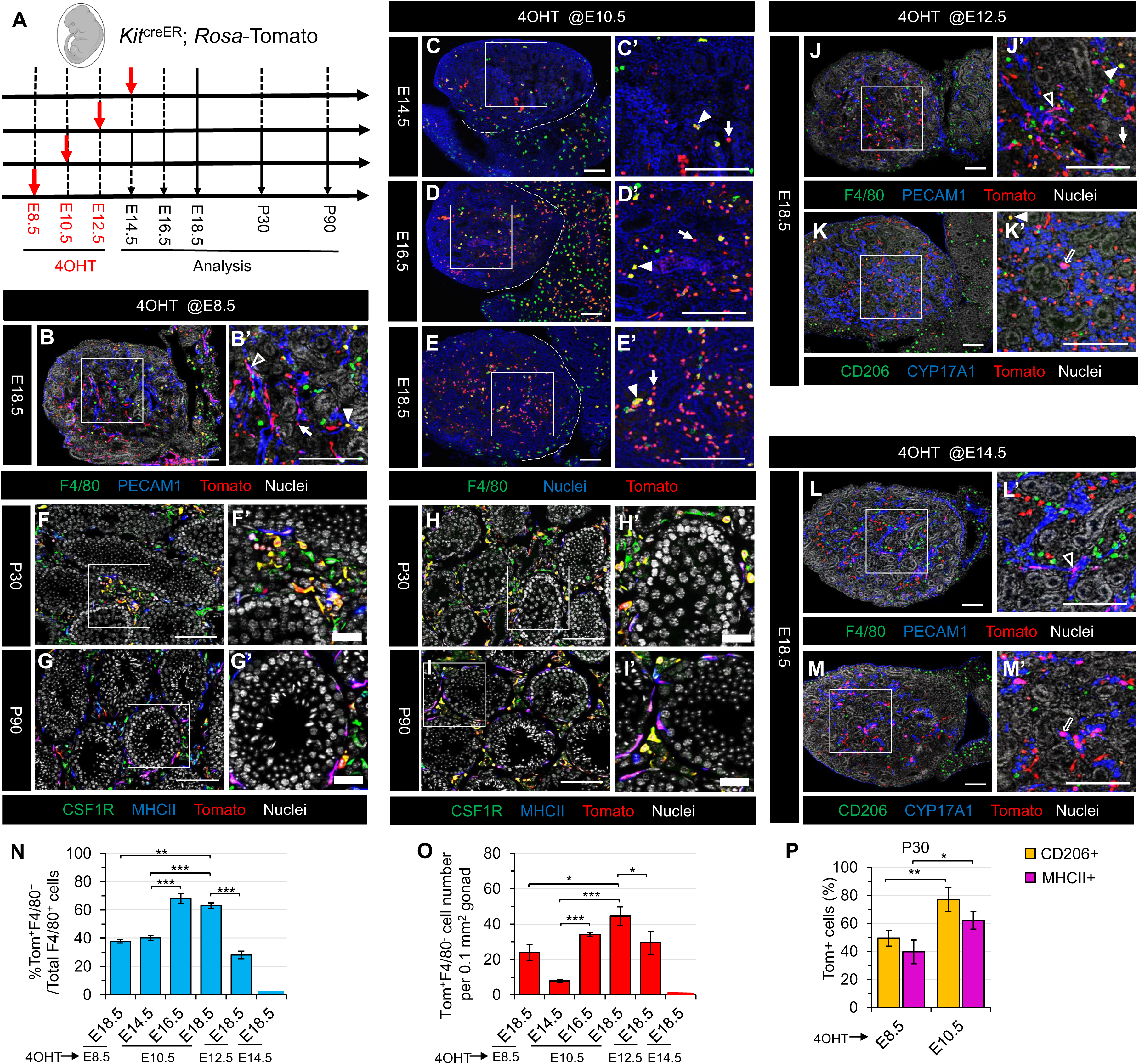
Contribution of AGM-derived HSCs to adult testicular macrophages occurs during a specific recruitment window during fetal development. (A) Strategy for 4OHT-induced lineage-tracing and harvesting of testes from *Kit*^creER^; *Rosa*-Tomato embryos and juvenile/adult mice. (B-N) Images of testes at various stages from *Kit*^creER^; *Rosa*-Tomato mice exposed to 4OHT at E8.5 (B,F,G), E10.5 (C-E,H,I), E12.5 (J,K), or E14.5 (L,M). Dashed lines indicate gonad-mesonephros boundary. Black arrowheads denote Tomato-expressing PECAM1+ endothelial cells, white arrowheads denote Tomato-expressing F4/80+ macrophages, black arrows denote Tomato-expressing CYP17A1+ Leydig cells, and white arrows denote Tomato-expressing F4/80-negative cells. Thin scale bar, 100 μm; thick scale bar, 25 μm. (N-P) Graphs showing quantification (*n*=3) of percent Tomato-expressing F4/80+ macrophages at E14.5, E16.5, or E18.5 Tomato-expressing interstitial (CD206+) and peritubular (MHCII+) macrophages at P30 (P) in *Kit*^creER^; *Rosa*-Tomato testes induced with 4OHT at various embryonic stages. Data are shown as mean +/- SD. \**P*<0.05; \*\**P*<0.01; \*\*\**P*<0.001 (two-tailed Student’s *t*-test).

When pulsed at E8.5, we detected Tomato^+^F4/80^+^ testicular macrophages and Tomato^+^F4/80^-^ cells in E18.5 testes (Fig. 2B, 2N and 2O), in contrast to the E8.5-induced-*Csf1r-*creER model, which had virtually no Tomato^+^F4/80^-^ cells. However, consistent with the *Csf1r-*creER results, microglia in E18.5 brains were labeled by Tomato (Fig. S4A), which is consistent with a previous study showing the labeling of AGM-derived multipotent progenitors (MPPs) (11) with an E8.5 pulse, suggesting that induction at E8.5 targets YS-derived EMPs and AGM-derived HSCs.

When pulsed at E10.5, we observed no Tomato-labeled microglia in E18.5 brains (Fig. S4B), while there was extensive labeling of macrophages, B cells and T cells in the postnatal liver and spleen (Fig. S4D-I), suggesting that induction at E10.5 can specifically label AGM-derived HSCs that give rise to all hematopoietic cell lineages. We detected a small number of Tomato+ cells in E14.5 testes when induced at E10.5 (Fig. 2C), but there was a considerable increase in the proportion of Tomato^+^F4/80^+^ macrophages and in the number of Tomato^+^F4/80^-^ cells from E16.5 to E18.5 when compared to induction at E8.5 (Fig. 2D, 2E, 2N and 2O). In E18.5 testes, all Tomato-expressing cells were CD45+ and divided into two populations, which were CD45^+^IBA1^+^ (IBA1 is a microglial/macrophage marker) and CD45^+^IBA1^-^ (Fig. S4C), suggesting that CD45^+^Tomato^+^F4/80^-^/ IBA1^-^ cells in E16.5 or E18.5 testes may represent monocytes. In addition, induction at E10.5 produced a higher number of Tomato+ interstitial and peritubular macrophages in P30 and P90 testes compared to induction at E8.5 (Fig. 2F-I). This may be due to the incomplete labeling of fetal HSCs when pulsed at E8.5. These results suggest that AGM-derived HSCs give rise to testicular interstitial and peritubular macrophages.

To further study whether fetal liver HSCs contribute to testicular macrophages, we induced Tomato expression in *Kit*^creER^; *Rosa-*Tomato embryos at E12.5 or E14.5 when the fetal liver has become the dominant site of hematopoiesis (13). In E18.5 testes that were induced at E12.5, there was a similar labeling efficiency of F4/80^+^ macrophages and a comparable Tomato^+^F4/80^-^ cell number as compared to induction at E8.5, but levels were relatively lower than with E10.5 induction (Fig. 2J, 2N and 2O). However, induction at E14.5 showed no Tomato-labeled testicular macrophages (Fig. 2L, 2N and 2O), suggesting that the contribution of fetal HSCs to testicular macrophages has a limited recruitment window. We also found that the number of Tomato-labeled CYP17A1+ fetal Leydig cells in E18.5 testes dramatically increased when induced at E14.5 as compared to E12.5 (Fig. 2K and 2M), which is linked to KIT expression in fetal Leydig cells (40, 41). Surprisingly, only a few PECAM1+ endothelial cells in E18.5 testis were labeled (Fig. 2J and 2L), unlike the broadly labeled endothelial cells that appeared in both testis and mesonephros when induced at E8.5, supporting the previous observation that YS-derived EMPs contribute endothelial cells to various tissue blood vessels (32).

To further investigate whether nascent and adult HSCs contribute to testicular macrophages, we exposed *Kit*^creER^; *Rosa-*Tomato pups and adult mice to tamoxifen (TAM) at P4 and P5, and P60-P62, respectively (Fig. S5). When injected at P4 and P5, we found that no testicular macrophages were labeled (Fig. S5A-C), whereas endothelial cells (PECAM1+), spermatogonia (KIT+) and Leydig cells (KIT+, CYP11A1+ or SF1+) expressed Tomato in P7, P30 and P60 testis (Fig. S5D-I). Consistent with induction at the neonatal stage, in adult-induced testes 60 days post-induction (P120) we also observed no Tomato-labeled interstitial or peritubular macrophages (Fig. S5J), but with abundant labeling of Leydig cells (Fig. S5K and S5L). These results suggest that bone marrow-derived HSCs after birth have no contribution to testicular macrophages.

### Flt3+ fetal HSC-derived multipotent progenitors give rise to testis macrophages

To further confirm the contribution of HSCs to testicular macrophages, we used *Flt3-*cre; *Rosa-*Tomato mice, in which Cre activity is mainly activated in late fetal or adult HSC-derived MPPs, but not in YS-derived EMPs and mature macrophages (7,42,43). We found that scattered Tomato+ cells appeared in E14.5 testes (Fig. 3A); however, there were numerous Tomato+ cells in E14.5 liver where extremely few of them were F4/80+ macrophages or KIT^+^CD45^-^ HSCs, but instead were either KIT^+^CD45^+^ or KIT^-^CD45^+^ progenitors (Fig. S6A and S6B), indicating that fetal liver hematopoietic progenitors do not migrate towards the testis at this stage. Subsequently, from E16.5 to P7, *Flt3-*cre; *Rosa-*Tomato testes exhibited extensive Tomato+ cells with a gradual increase in the percentage of F4/80^+^ testicular macrophages expressing Tomato in those stages as compared to E14.5 (Fig. 3B-D and 3I). As negative controls, in E18.5 liver and brain, less than 10% of macrophages expressed Tomato (Fig. S6C, S6D and S6G), consistent with a previous study (7), suggesting that liver and brain macrophages have reached a steady-state equilibrium at early fetal stages. However, Tomato-expressing cells were CD45^+^KIT^+^ or CD45^+^KIT^-^ fetal HSC-derived progenitors in E18.5 liver, and CD45+ myeloid cells in E18.5 testes (Fig. S6E and S6F). Additionally, the number of Tomato^-^F4/80^+^ cells during late fetal testis development also increased and plateaued in E18.5 testes, but then declined in P7 testes (Fig. 3B-D and 3J), implying that Tomato^+^F4/80^-^ cells in fetal testis may be the precursors of F4/80+ macrophages and differentiate into mature macrophages after birth. Indeed, in P30 and P90 testes, we observed an extremely high labeling efficiency of interstitial and peritubular testicular macrophages (Fig. 3E-H). These results suggest that fetal HSC-derived *Flt3*+ progenitors give rise to testicular macrophages.

**Figure 3.**
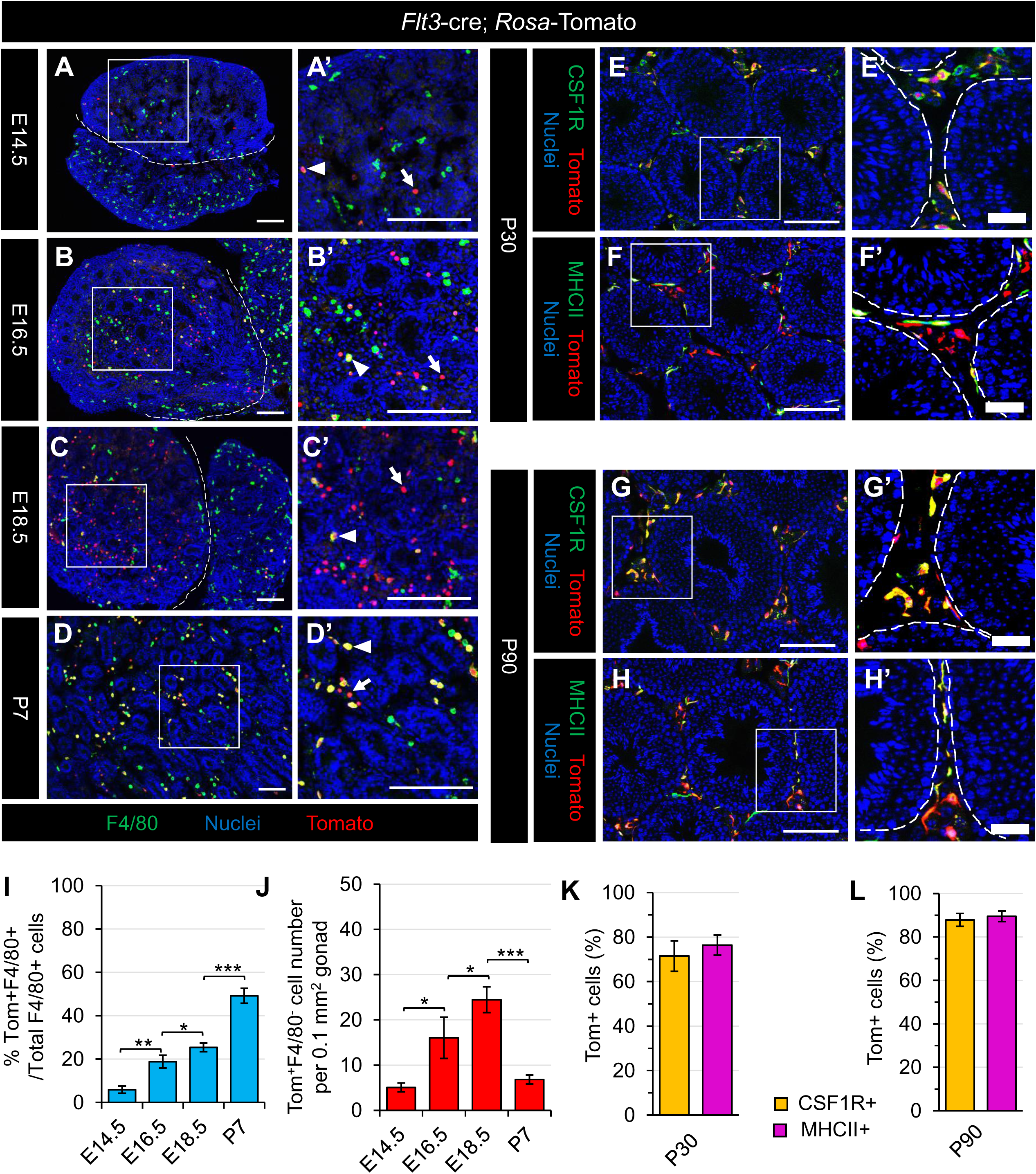
*Flt3*-expressing fetal HSC-derived multipotent progenitors give rise to testicular macrophages. (A-H) Images of *Flt3*-cre; *Rosa*-Tomato testes at E14.5 (A), E16.5 (B), E18.5 (C), P7 (D), P30 (E-F), and P90 (G-H). Arrowheads denote Tomato-expressing F4/80+ macrophages and arrows denote Tomato-expressing F4/80-negative cells. Thin scale bar, 100 μm; thick scale bar, 25 μm. (I-L) Graphs showing quantification of percent Tomato-expressing F4/80+ macrophages (I), number of Tomato- expressing F4/80-negative cells per unit area at E14.5, E16.5, E18.5 or P7 (J), and percent Tomato- expressing interstitial (CD206+) and peritubular (MHCII+) macrophages at P30 (K) and P90 (L) in *Flt3-*cre; *Rosa*-Tomato testes. Data are shown as mean +/- SD. \**P*<0.05; \*\**P*<0.01; \*\*\**P*<0.001 (two-tailed Student’s *t*-test).

### Fetal testicular monocytes gradually differentiate into testicular macrophages after birth

Based on the above results showing the emergence of extensive “monocyte-like” cells in late fetal testes, we next attempted to establish whether fetal monocytes contribute to adult testicular macrophages. Although no CreER-based fate-mapping model can specifically track fetal liver-derived monocytes, the *Cx3cr1*^creER^ mouse model has been widely used in the fate-mapping of myeloid cells (12). *Cx3cr1* is mainly expressed in mononuclear phagocytes, including macrophages and monocytes, particularly Ly6C-negative or Ly6C- low monocytes (44, 45). We generated *Cx3cr1*^creER^; *Rosa-*Tomato mice, which were induced with 4OHT at E12.5, E18.5, and with tamoxifen (TAM) at postnatal stages (Fig. 4A).

**Figure 4.**
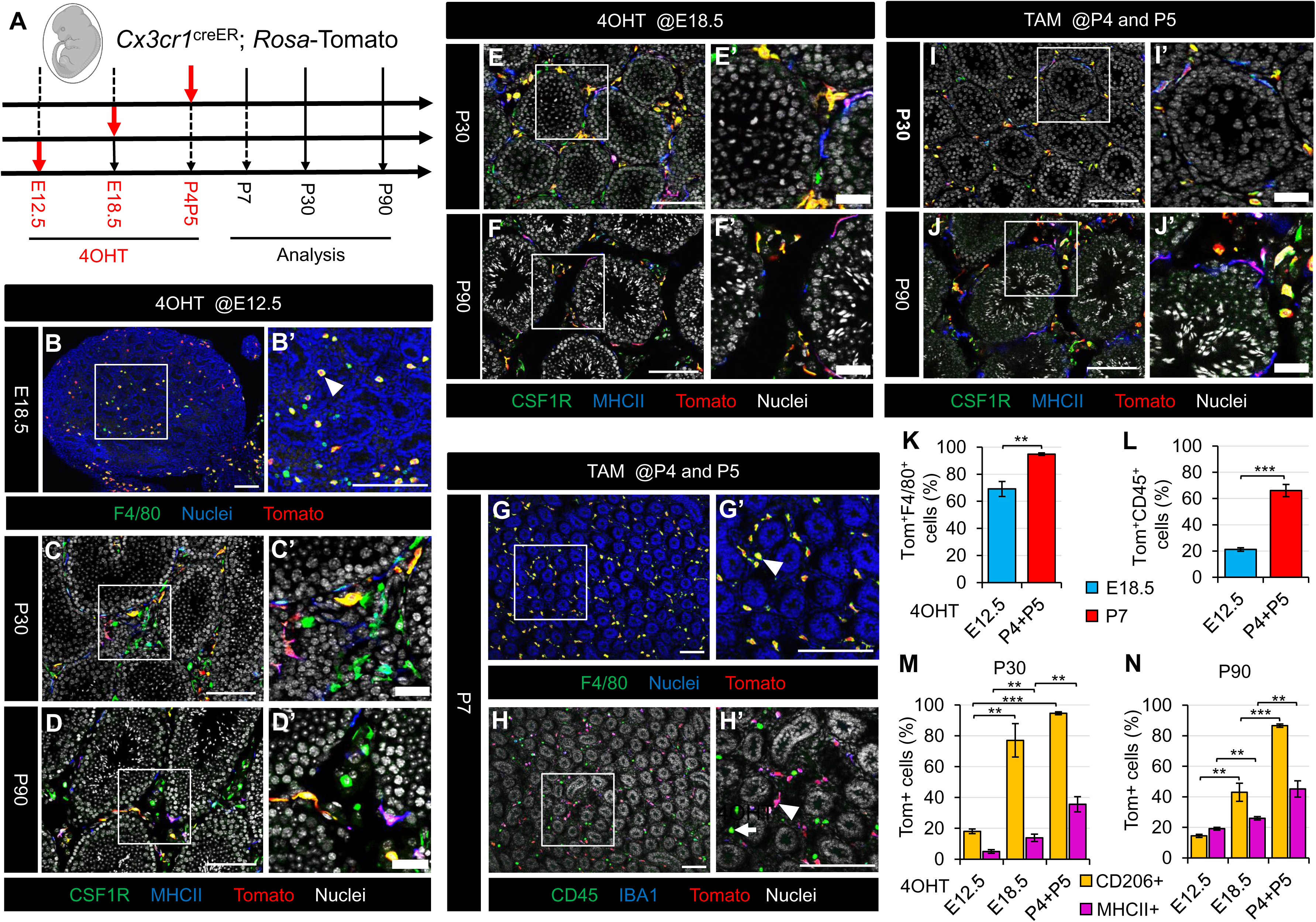
Fetal testis monocytes gradually differentiate into testicular macrophages after birth. (A) Strategy for 4OHT-induced lineage-tracing and harvesting of testes from *Cx3cr1*^creER^; *Rosa*-Tomato embryos and juvenile/adult mice. (B-J) Images of testes at various stages from *Cx3cr1*^creER^; *Rosa*-Tomato mice exposed to 4OHT at E12.5 (B-D), E18.5 (E-F), or P4 and P5 (G-J). Arrowheads denote Tomato-expressing IBA1+ or F4/80+ macrophages; arrow denotes a Tomato-negative CD45+ cell. Thin scale bar, 100 μm; thick scale bar, 25 μm. (K-N) Graphs showing quantification (*n*=3) of percent Tomato-expressing F4/80+ macrophages at E18.5 or P7 (K), percent Tomato-expressing CD45+ cells at E18.5 or P7 (L), and percent Tomato-expressing interstitial (CD206+) and peritubular (MHCII+) macrophages at P30 (M) or P90 (N) in *Cx3cr1*^creER^; *Rosa*-Tomato testes induced with 4OHT at various embryonic or postnatal stages. Data are shown as mean +/- SD. \*\**P*<0.01; \*\*\**P*<0.001 (two-tailed Student’s *t*-test).

When pulsed at E12.5, we observed that more than 60% of F4/80+ macrophages were labeled with Tomato in E18.5 testes (Fig. 4B and 4K), but few F4/80-negative cells were labeled, which resulted in a low percentage of Tomato labeling among total CD45^+^ cells (Fig. 4B and 4L), implying that *Cx3cr1-*CreER activity in early fetal liver-derived monocytes was inefficiently activated. In P30 and P90 testes, there were few Tomato+ interstitial and peritubular macrophages (Fig. 4C, 4D, 4M and 4N), suggesting that E12.5- induced testicular macrophages are severely diluted after birth in the adult. Given the presence of monocytes in E18.5 testes, we speculated that monocytes during the perinatal period may contribute to adult testicular macrophages. To test this hypothesis, we administered 4OHT to *Cx3cr1*^creER^; *Rosa-*Tomato embryos at E18.5. As expected, induction at E18.5 had relatively higher labeling of interstitial and peritubular macrophages in P30 and P90 testes as compared to induction at E12.5 (Fig. 4E, 4F, 4M and 4N), but also had a reduction in the percentage of Tomato-labeled interstitial macrophages with age increase (Fig. 4M and 4N), indicating a limited labeling of monocytes in E18.5 testes. When injected at P4 and P5, we found that more than 90% of F4/80+ macrophages expressed Tomato in P7 testes (Fig. 4G and 4K). Additionally, there was a significant increase in the percentage of Tomato-labeled testicular interstitial and peritubular macrophages at P30 and P90 (Fig. 4I, 4J, 4M and 4N), with interstitial macrophage labeling remaining steady (Fig. 4M and 4N). This may be due to the increased recombinase efficiency of CreER activated from E12.5 as compared to neonatal stages, with the gradual upregulation of CX3CR1 expression and downregulation of Ly6C expression in fetal monocytes when they infiltrate from the fetal liver to their target tissue (8). We also found a large number of unlabeled CD45^+^IBA1^-^ monocytes in P7 testes (Fig. 4H and 4L), which were responsible for the low labeling efficiency of peritubular macrophages. These results suggest that fetal monocytes that colonize late fetal testes contribute to both adult testicular macrophage populations.

### Sertoli cells instead of germ cells regulate macrophage recruitment and differentiation

We next assessed which cell types in the testis microenvironment, such as germ cells or Sertoli cells, are important for fetal monocyte recruitment and differentiation into adult testicular macrophages. To test whether Sertoli cells contribute to recruitment of fetal testicular monocytes, we ablated Sertoli cells specifically by activating the expression of diphtheria toxin fragment A (DTA) in *Rosa-*DTA mice by crossing with *Amh-*cre mice, in which Cre recombinase is exclusively expressed in Sertoli cells (46). We found that in *Amh-*cre; *Rosa-*DTA testes from E14.5 to E18.5, a majority of testis cords with SOX9+ or AMH+ Sertoli cells were ablated, with only a few testis cords near the gonad-mesonephros border region remaining (Fig. 5A-L), consistent with previous studies on the effectiveness of Sertoli cell depletion in fetal *Amh-*cre; *Rosa-*DTA testes (47, 48). Although F4/80+ macrophage number in Sertoli cell-depleted testes was comparable to control testes from E14.5 to E18.5 (Fig. 5A-F and 5M), there was a significant reduction in the number of CD45^+^IBA1^-^ monocytes in Sertoli cell-depleted testes from E16.5 to E18.5 (Fig. 5G-L and 5N). These results suggest that Sertoli cells play a major role in fetal testicular monocyte recruitment. We also observed a decrease in the number of blood vessels in Sertoli cell-depleted testes (Fig. 5A-L), which may impede the migration of fetal monocytes from the fetal liver to the testis.

**Figure 5.**
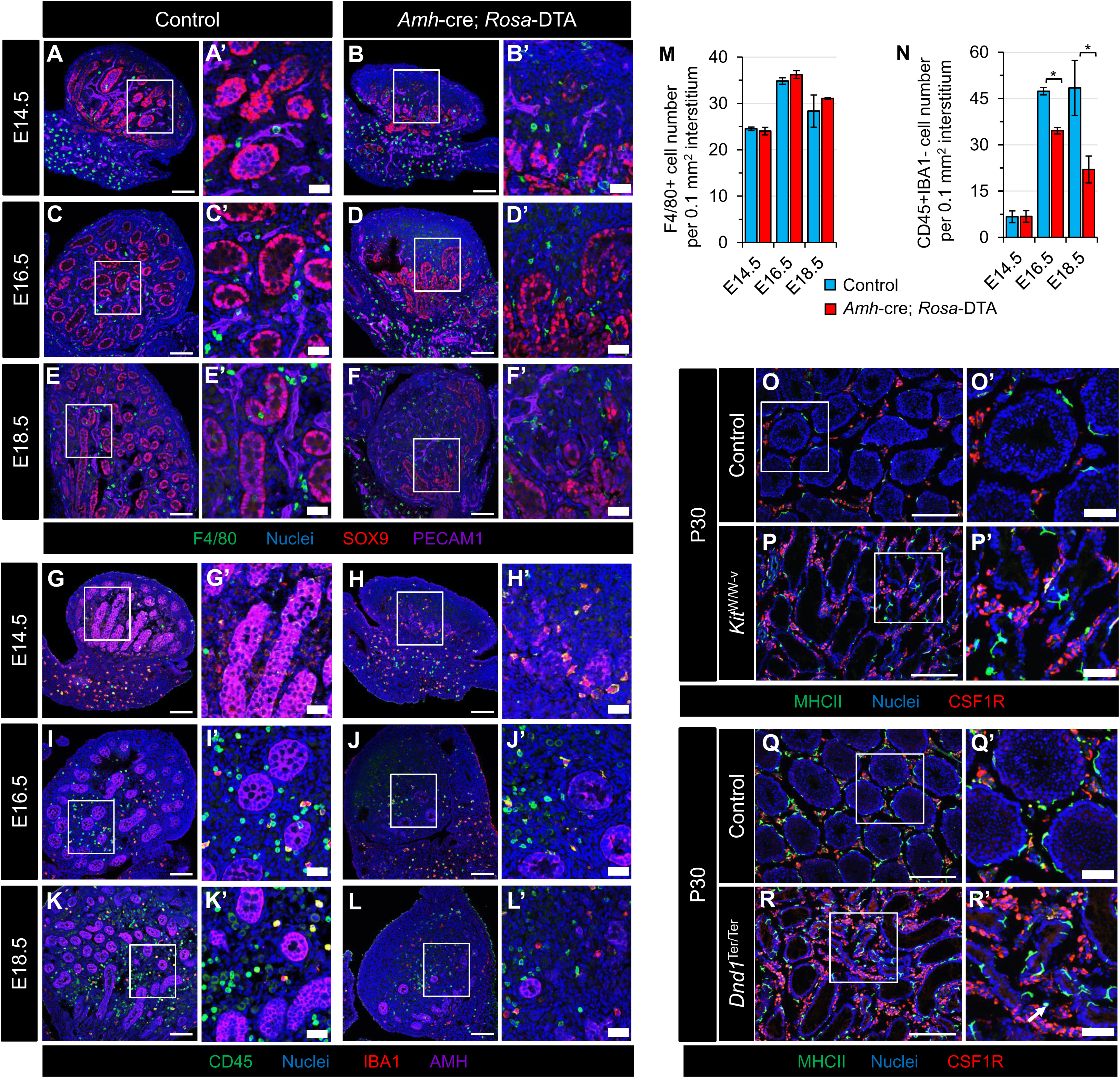
Sertoli cells, not germ cells, regulate testicular macrophage recruitment and differentiation. (A-L) Images of control (A,C,E,G,I,K) and *Amh-*cre; *Rosa*-DTA (B,D,F,H,J,L) testes at various fetal stages. (M-N) Graphs showing number of F4/80+ cells (M) or number of CD45+ IBA1-negative cells (N=3) per unit area at E14.5, E16.5, or E18.5 in control versus *Amh-*cre; *Rosa*-DTA fetal testes. Data are shown as mean +/- SD. \**P*<0.05 (two-tailed Student’s *t*-test). (O-P) Images of P30 control (O and Q), *Kit^W/W-v^* mutant (P), and *Dnd1^Ter/Ter^* mutant (R) testes. Arrow denotes abnormal interstitially-localized MHCII+ macrophage in the *Dnd1^Ter/Ter^* mutant testis. Thin scale bar, 100 μm; thick scale bar, 25 μm.

To further examine the role of postnatal Sertoli cells on macrophage differentiation, we analyzed testicular macrophage populations in *Dmrt1*-null knockout (*Dmrt1*^-/-^) males (49), which exhibit a postnatal loss of Sertoli cell identity and the transdifferentiation of Sertoli cells to granulosa cells (50, 51), which is the somatic supporting cell type of the ovarian follicle. We found that during postnatal *Dmrt1*^-/-^ testis development, SOX9+ Sertoli cells gradually transdifferentiated into FOXL2+ granulosa cells with an apparent increase in F4/80+ macrophage density (Fig. S7A-F), likely partly due to smaller testis size. In P7, P18 and P60 *Dmrt1*^-/-^ testes, CSF1R+ interstitial macrophages were still located in the interstitial compartment similar to *Dmrt1*^+/-^ control testes (Fig. S7G-L). Because MHCII+ peritubular macrophages only emerge postnatally after 2 weeks of age (23), *Dmrt1*^-/-^ testes had normal seminiferous tubules and similar MHCII+ round cells, similar to P7 control testes (Fig. S7G and S7H). At P18, unlike control testes that showed normal MHCII+ peritubular macrophage distribution around seminiferous tubules, many MHCII+ round cells were observed in the interstitial compartment of P18 *Dmrt1*^-/-^ testes (Fig. S7I and S7J). By P60, MHCII+ round cells had developed into macrophages with a stellate or dendritic morphology, but were still retained in the interstitial compartment and were not properly localized to seminiferous tubule surfaces (Fig. S7K and S7L). These results suggest that postnatal Sertoli cells are critical for peritubular macrophage differentiation and localization.

Given the germ cell loss in *Amh-*cre; *Rosa-*DTA and *Dmrt1* knockout testes (47, 51), we next explored whether germ cells have an effect on macrophage differentiation. We used *Kit*^W/W-v^ and *Dnd1^Ter/Ter^* mouse models, both of which show germ cell deficiency, but have otherwise normal seminiferous tubules (52, 53). In P30 *Kit^W/W-v^* and *Dnd1^Ter/Ter^* testes, we observed normal localization and morphology of CSF1R+ interstitial and MHCII+ peritubular macrophages as compared to control testes (Fig. 5O-R), suggesting that germ cells have no significant role in macrophage differentiation. Overall, our results indicate that Sertoli cells, rather than germ cells, promote macrophage differentiation after birth.

### EMPs and HSC-derived macrophages have distinct functions in fetal testis development

To determine the specific function(s) of distinct fetal testicular macrophage populations during embryogenesis, we first need to verify which hematopoietic waves contribute to early fetal testicular macrophages. Therefore, we exposed *Csf1r-*creER; *Rosa-*Tomato and *Kit*^creER^; *Rosa-*Tomato embryos to 4OHT at E8.5 and E10.5 to label YS-derived EMPs and AGM-derived HSCs, respectively. In *Csf1r-*creER; *Rosa-*Tomato embryos, there were extensive Tomato-labeled F4/80+ macrophages in E12.5 gonad/mesonephros complexes induced at E8.5 or E10.5 (Fig. S8A and S8B), whereas CD11b+ monocytes located in the gonad-mesonephros border region were labeled only after induction at E8.5 (Fig. S8C and S8D). In *Kit*^creER^; *Rosa-*Tomato embryos, some F4/80+ macrophages and CD11b+ monocytes in E12.5 gonad/mesonephros complexes were Tomato-positive when induced at E8.5, instead of at E10.5 (Fig. S8E- H). In addition, some DDX4+ (also called MVH) germ cells were labeled by Tomato in both E8.5- and E10.5-induced gonads (Fig. S8G and S8H), consistent with KIT expression in primordial germ cells (54). These lineage-tracing results demonstrate that early fetal testicular macrophages originate from YS-derived EMPs.

Since the development and migration of YS-derived macrophage progenitors are dependent on CSF1R (6), we next attempted to deplete YS macrophage progenitors by transiently inhibiting the CSF1R signaling pathway in embryos injected with a blocking anti-CSF1R antibody at E6.5, as reported previously (8, 55). CSF1R antibody injection resulted in efficient depletion of F4/80+ macrophages in E13.5 testes (Fig. 6A and 6B); however, we observed an increase in the number of CD11b+ monocytes concentrated in the gonad-mesonephros border region (Fig. 6C and 6D). qRT-PCR analyses also demonstrated a significant downregulation of macrophage-related gene expression, such as *Cx3cr1*, *Csf1r*, and *Adgre1* (also called *F4/80*) in CSF1R-antibody-injected E13.5 gonads compared to controls (Fig. 6E). Despite no change in the expression of *Sox9* and *Amh* (two Sertoli cell genes) (Fig. 6E), morphometric analyses of CSF1R-antibody-injected E13.5 gonads showed a reduction in testis cord width but no effect on testis cord height (Fig. 6F and 6G). Furthermore, the percentage of abnormal, non-linear testis cords, including fused and branched cords, as well as cord bridges, was increased in CSF1R-antibody-injected E13.5 gonads (Fig. 6H). These data demonstrate the impact of distorted myeloid cell ratios is specific to testis cord morphogenesis rather than Sertoli cell gene expression. To assess if macrophage depletion had any effect on the development of other cell types, including endothelial cells, Leydig cells, and germ cells in E13.5 gonads, we assessed the expression of endothelial cell (*Cdh5*), Leydig cell (*Cyp11a1*, *Cyp17a1* and *Hsd3b1*), and germ cell (*Kit*; *Pou5f1*, also called *Oct4*; and *Ddx4*) genes. However, they were no significant or detectable differences in expression based on qRT-PCR and immunofluorescent analyses (Fig. 6E).

**Figure 6.**
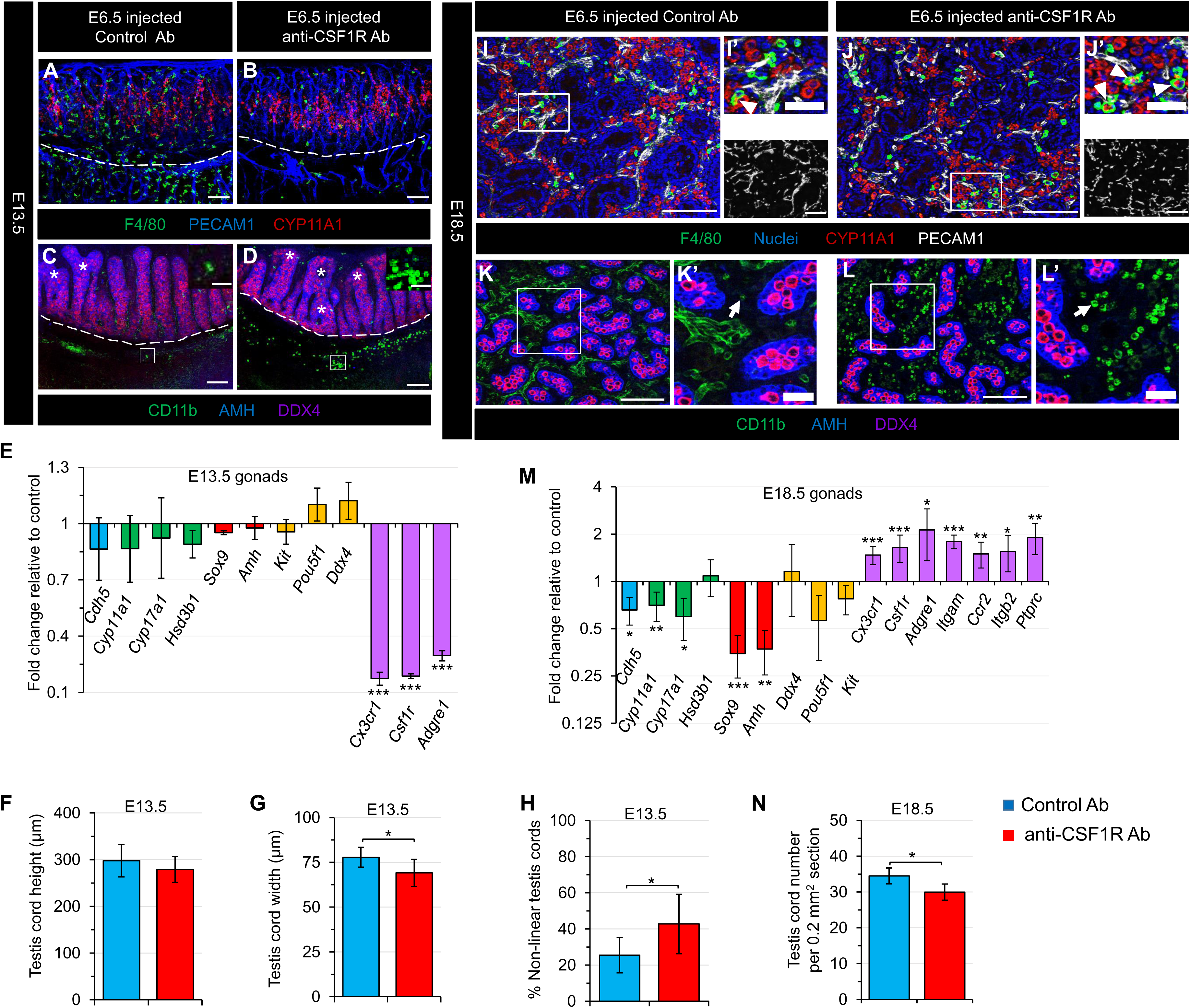
EMPs and HSC-derived macrophages have distinct functions during fetal testis development. (A-D) Images of E13.5 fetal testes from C57BL/6J embryos exposed at E6.5 to either control rat IgG2a antibody (A,C) or anti-CSF1R blocking antibody to deplete YS-derived macrophages (B,D). Dashed lines indicate gonad-mesonephros boundary. Asterisks denote fused or branched testis cords. Insets in C and D are higher-magnification images of the boxed regions highlighting CD11b+ monocytes in the gonad-mesonephros border region. (E) qRT-PCR analyses of whole E13.5 fetal testes showing fold change of expression in YS-macrophage-depleted samples versus controls. Graph data are shown as mean +/- SD. \*\*\**P*<0.001 (two-tailed Student’s *t*-test). (F-H) Graphs showing quantification of testis cord height (F), testis cord width (G), and percent abnormal (fused and/or branched) testis cords (H) in YS-macrophage-depleted samples versus controls. (I-L) Images of E18.5 fetal testes from C57BL/6J embryos exposed at E6.5 to either control rat IgG2a antibody (I,K) or anti-CSF1R blocking antibody to deplete YS-derived macrophages (J,L). Arrowheads denote cells exhibiting co-expression of F4/80 and CYP11A1; arrows denote CD11b+ monocytes. (M) qRT-PCR analyses of whole E18.5 fetal testes showing fold change of expression in YS- macrophage-depleted samples versus controls. (N) Graph showing testis cord number per unit area in YS- macrophage-depleted samples versus controls. Thin scale bar, 100 μm; thick scale bar, 25 μm. All graph data are shown as mean +/- SD. \**P*<0.05; \*\**P*<0.01; \*\*\**P*<0.001 (two-tailed Student’s *t*-test).

We found that testicular macrophages in CSF1R-antibody-exposed embryos were fully repopulated by E18.5 (Fig. 6I and 6J), with extensive CD11b+ monocytes (Fig. 6K and 6L), suggesting the migration of monocytes in the E13.5 gonad-mesonephros border region into the gonad. Additionally, the mRNA expressions of macrophage-related genes, such as *Cx3cr1*, *Csf1r*, and *Adgre1,* and monocyte-related genes, such as *Itgam* (also called CD11b), *Itgb2* (also called CD18), *Ccr2* and *Ptprc* (also called CD45) were significantly increased in CSF1R-antibody-injected E18.5 testes compared to controls (Fig. 6M). These results suggest that YS-derived EMPs are dispensable for giving rise to late fetal testicular macrophages, and that alternative embryonic precursors could functionally replace early fetal testis macrophages originating from YS-derived EMPs during development, supporting our above-demonstrated findings. We observed more CYP17A1+ Leydig cells colocalized with macrophages in CSF1R-antibody-injected E18.5 testes (Fig. 6I and 6J), perhaps indicative of phagocytosis, with a reduction in mRNA levels of the Leydig cell genes *Cyp11a1*and *Cyp17a1* (Fig. 6M), suggesting an ectopic engulfment role of repopulated macrophages. Except for Leydig cells, gonads with repopulated macrophages also had reduced expression of *Cdh5*, *Sox9* and *Amh* (Fig. 6M), along with more intermittent blood vessels (Fig. 6I and 6J) and a reduction in the number of AMH+ testis cords (Fig. 6K and 6L, 6N). However, mRNA levels of germ cell genes were not changed (Fig. 6M). These data suggest that repopulated excess testicular macrophages after prior CSF1R-antibody-mediated depletion have adverse effects on multiple cellular niches in late fetal testes.

### Adult testicular macrophages promote Leydig cell proliferation and steroidogenesis

Given adult interstitial macrophages located within the testicular interstitium are in close contact with Leydig cells (26, 29), we hypothesized that interstitial macrophages regulate adult Leydig cell development and steroidogenesis. To address this hypothesis, we used anti-CD45 and anti-F4/80 magnetic antibody microbeads to remove or enrich macrophages from isolated primary testicular interstitial cells in vitro via magnetic-activated cell sorting (MACS). We first assessed the efficiency of testicular macrophage depletion and enrichment in interstitial cells from adult *Cx3cr1*^GFP^ testes, in which testicular macrophages express GFP (26). We found that GFP+ testicular macrophages were nearly completely depleted in the CD45- fraction and abundantly enriched in the CD45+ and F4/80+ fractions, particularly the F4/80+ fraction (Fig. S9A), as compared to pre-separation interstitial cell populations. Additionally, about 90% of GFP+ cells expressed CD206 (Fig. S9A), suggesting that macrophages enriched by anti-CD45 or anti-F4/80 microbeads were testicular interstitial macrophages.

We then cultured the pre-separation, CD45-, CD45+ and F4/80+ interstitial cell populations, using different ratios of testicular macrophages and Leydig cells as a result of the MACS purification, in vitro for 3 days and 6 days. Flow cytometry results showed that in the CD45- fraction, almost no CD206+ testicular macrophages appeared and most of them were CD106+ (encoding VCAM1), a Leydig cell marker (56), especially on day 6 (Fig. S9B and S9C). Surprisingly, the percentage of CD206+ testicular macrophages on day 3 in in the CD45+ and F4/80+ fractions was lower than at the onset of culture, but by day 6 it returned to high levels (Fig. S9B and S9C), indicating a varied proliferative capacity of testicular macrophages and Leydig cells at different time points during co-culture. To verify this potential trend, we labeled dividing cells using an EdU incorporation assay. Immunofluorescence analysis revealed a higher percentage of EdU^+^CYP11A1^+^ proliferating Leydig cells in all fractions on day 3 than day 6 (Fig. 7A-I), suggesting that Leydig cell proliferative potential diminished as culture time in vitro was extended. In addition, on day 3 rather than day 6, there was a significant increase in the percentage of EdU+ Leydig cells in the CD45+ and F4/80+ fractions compared to the pre-separation fraction, with an even greater increase when compared to the CD45- fraction (Fig. 7A-I), suggesting the initial proliferation of Leydig cells is dependent on testicular macrophages. However, CD206+ testicular macrophages had a stronger proliferative capacity in the CD45+ and F4/80+ fractions on day 6 (Fig. 7A-H, 7J).

**Figure 7.**
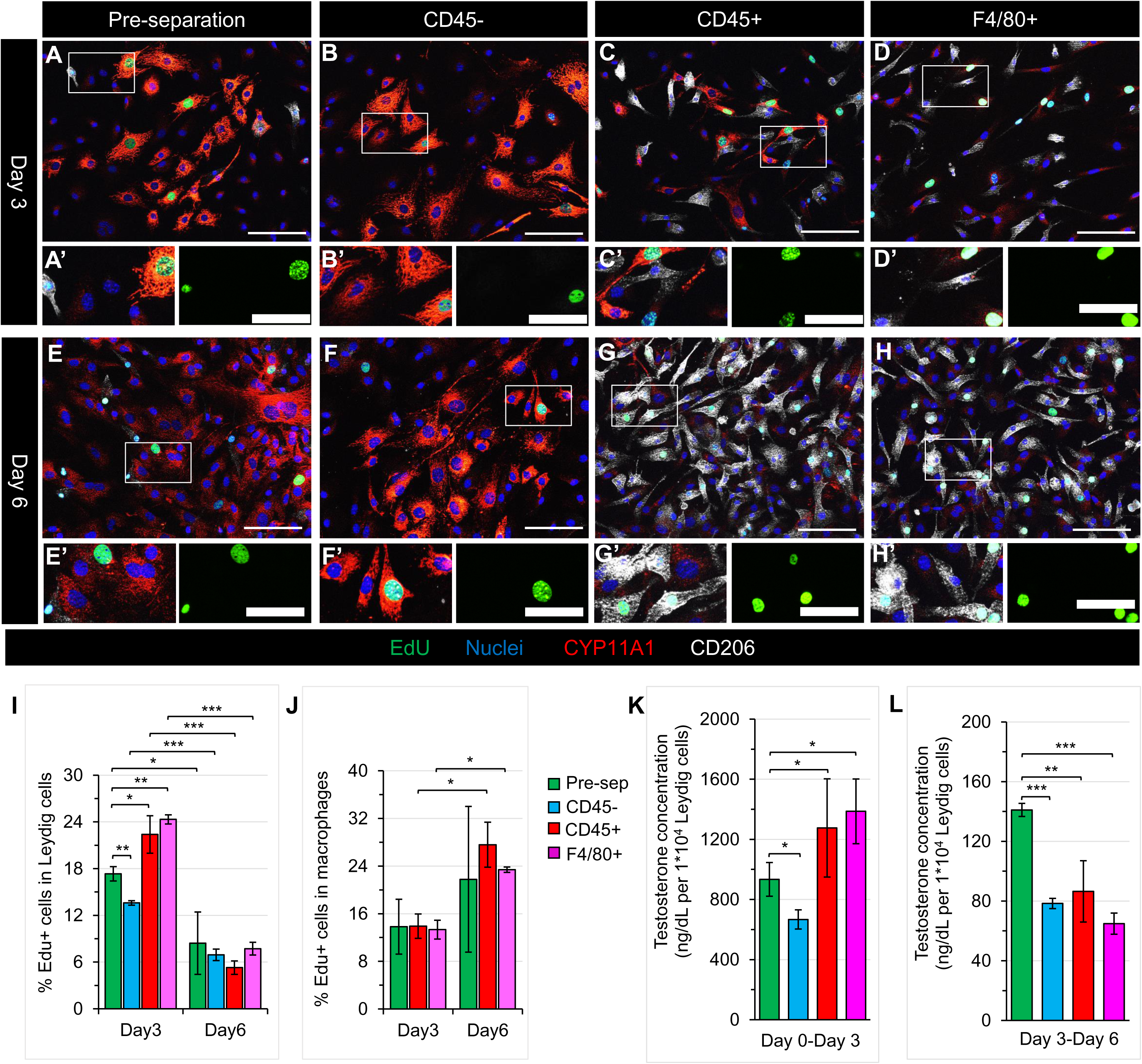
Adult testicular macrophages promote Leydig cell proliferation and steroidogenesis. (A-H) Images of primary cell culture after 3 days (A-D) and 6 days (E-H) from adult (3-month-old) C57BL/6J testes for pre-separation (A,E), CD45-depleted (B,F), CD45-enriched (C,G), and F4/80-enriched (D,H) populations. (I-L) Graphs showing percent EdU+ Leydig cells (I), percent EdU+ macrophages (J), testosterone concentration in culture media after 0-3 days of culture (K), and testosterone concentration in culture media after 3-6 days of culture (L) in the 4 different cell populations. Thin scale bar, 100 μm; thick scale bar, 25 μm. All graph data are shown as mean +/- SD. \**P*<0.05; \*\**P*<0.01; \*\*\**P*<0.001 (two-tailed Student’s *t*-test).

To define the role of testicular macrophages in Leydig cell steroidogenesis, we measured testosterone levels in the supernatant of each interstitial cell fraction on day 3 and day 6 of culture. We found testosterone concentrations were significantly reduced in the supernatant of the CD45- fraction, and were increased in the supernatant of the CD45+ and F4/80+ fractions compared to the pre-separation fraction from day 0 to day 3 (Fig. 7K). However, from day 3 to day 6, Leydig cells produced extremely low levels of testosterone in all fractions, with even lower testosterone levels in the CD45-, CD45+, and F4/80 fractions, which contained either no macrophages or extensive macrophages (Fig. 7L). These data suggest the proper number of testicular macrophages promotes Leydig cell steroidogenesis, but that an excess of testicular macrophages may be detrimental to this process.

## Discussion

Although it is clear that there are at least two distinct macrophage populations in the adult testis (22,23,26), there has been controversy in recent studies about the hematopoietic origins of testicular macrophages (22–24). Here our analysis of multiple lineage-tracing models definitively demonstrated that in late gestation, fetal testicular monocytes originating from AGM-derived fetal HSCs give rise to the two adult testicular macrophage subsets, CSF1R^+^CD206^+^MHCII^-^ interstitial macrophages and CSF1R^-^CD206^-^MHCII^+^ peritubular macrophages, and that YS-derived EMPs have a minor contribution. However, HSCs arising from either the neonatal or adult bone marrow do not contribute to adult testicular macrophages, at least in a normal developmental context. These findings emphasize the critical contribution of AGM-derived fetal HSCs during a limited developmental window to late fetal and adult testicular macrophage populations and address a knowledge gap in our understanding of testis-resident macrophage ontogeny.

Previous studies described two distinct macrophage populations in E14.5 to E18.5 testes, CD11b^low^F4/80^hi^ (which represent mature macrophages) and CD11b^hi^F4/80^low^ (which represent monocytes), the latter with a gradual increase in its representation within the gonad (22, 24), similar with our immunofluorescent results showing the co-existence of CD45^+^F4/80^+^ and CD45^+^F4/80^-^ immune cell types. However, the embryonic origins of late fetal testicular monocytes are still unknown. It is well-known that *Csf1r*-creER lineage tracing could unambiguously and specifically trace YS-derived EMPs when induced at E8.5 (7, 32), while it traced both HSC and YS-derived *Csf1r*+ myeloid cells when pulsed at E10.5 and E12.5 in our study and in a recent study (33). In *Kit*^creER^ fate-mapping analysis, our results definitively showed that it could specifically trace AGM-derived HSCs in E10.5-induced embryos. Although a previous report claimed that the *Flt3*-cre approach robustly marks adult HSC progeny and has relatively weak labeling of fetal HSC output (7), our results showed extensive Kit*^-^*Flt3^+^CD45^+^progenitors in fetal liver starting from E14.5, suggesting the feasibility of using *Flt3*-cre mice for targeting or labeling fetal HSC-derived multipotent progenitors. By combining these lineage-tracing models, we found that a large number of CD45^+^F4/80^-^ monocytes infiltrate into the fetal testis starting from E16.5, in which they are derived from fetal HSCs, instead of YS-derived EMPs. Our previous studies showed that YS-derived primitive progenitors give rise to the early gonadal– mesonephric macrophages (21). In this study, we also observed the contribution of YS-derived EMPs to early fetal testicular macrophages, but a decrease in E18.5 testes, suggesting an influx of new macrophages. Additionally, when YS progenitors are depleted, the full recovery of F4/80+ macrophages in E18.5 testes from macrophage-depleted early gonads verified their non-YS origin. Indeed, our fate-mapping experiments revealed that fetal HSCs give rise to late fetal testicular F4/80+ macrophages, suggesting that a subset of fetal HSC-derived monocytes have differentiated into testicular macrophages during development. Therefore, our results demonstrated a dominant contribution of fetal HSCs to late fetal testicular macrophages and monocytes.

Except for microglia, which represent the sole tissue-resident macrophage population generated entirely from YS progenitors, most tissue-resident macrophages in steady state are derived from fetal HSCs (11). Our long-term lineage-tracing analyses showed that testicular interstitial and peritubular macrophages in adult mice are also mainly derived from fetal HSCs, supporting the “fetal HSC hypothesis” (11) and argue against the “EMP hypothesis” claiming that YS-derived EMPs are the main source of adult tissue-resident macrophages (8). In contrast, Mossadegh-Keller *et al.* (23) transferred adult bone-marrow-derived HSCs intrahepatically into neonatal pups to address the hypothesis that the peritubular macrophages are derived from bone marrow progenitors, while the early-embryo-derived testicular interstitial macrophages are partially replaced by replenishment from bone-marrow-derived cells. However, our *Kit*^creER^ lineage-tracing model induced at the adult stage showed that bone-marrow-derived HSCs make no contribution to testicular macrophages. *Ccr2*^-/-^ and parabiosis experiments from Lokka *et al.*(22) and Wang *et al.*(24) demonstrated the CCR2-independent characteristics of testicular macrophage populations, despite the fact that *Ccr2*^-/-^ mice exhibit a substantial loss of bone-marrow-derived monocytes in numerous postnatal organs and blood. The discrepancy between the findings of recent studies and a previous study by Mossadegh-Keller et al. may be due to a systemic inflammation after bone marrow cell injection (57) resulting in the contribution of bone marrow-derived cells to peritubular macrophages under inflammatory conditions (24), rather than in a normal physiological process. In addition, as a negative control, microglia only derived from YS progenitors are not investigated in HSC transplantation experiments. Although Lokka *et al.*(22) claimed that fetal-liver-derived monocytes contribute to both testicular macrophage populations using a *Cx3cr1*^creER^ mouse model injected with 4OHT at E13.5, the frequencies are exceedingly low, as shown by our results. However, when induced at the perinatal stage, our findings revealed that fetal testicular monocytes have a significant contribution to adult testicular macrophages. Thus, the *Cx3cr1*^creER^ mouse model is suboptimal for tracking early fetal-liver-derived monocytes, but its labeling efficiency could improve later in development due to a phenotype shift of Ly6C^hi^CX3CR1^lo^ toward Ly6C^lo^CX3CR1^hi^ when monocytes from the fetal liver migrate into target organs (12, 58). Wang *et al*. (24) claimed a neonatal bone-marrow-derived monocyte contribution to testicular macrophages by analyzing the frequency of *Flt3*-cre+ cells only in the postnatal testes of *Flt3-* cre mice. However, our results showed that *Flt3*-cre+ monocytes appear in late fetal testes. Additionally, our *Kit*^creER^ fate-mapping analyses revealed that neonatal HSCs have no contribution to adult testicular macrophages, contradicting a previous study’s conclusion (24). Altogether, our data definitively demonstrate that a small subset of adult testicular macrophage populations arise from YS-derived EMPs, but the vast majority are derived from fetal HSC-derived monocytes that arrive after YS-derived cells, and not from postnatal HSC-derived cells.

Growing evidence suggests that Sertoli cells modulate the immune privilege of the testis by forming the blood-testis-barrier (BTB)/Sertoli cell barrier, which prevents immune cells from obtaining access to and sequesters potentially auto-antigenic post-meiotic germ cells (16, 59). Furthermore, Sertoli cells can induce the recruitment of regulatory T cells (Tregs) from naïve T cells which, in turn, promotes Sertoli cell immunosuppressive functions in graft survival (60). In our study, we demonstrated that Sertoli cells are essential for fetal testicular monocyte recruitment by using an effective Sertoli-cell-specific cell ablation model: a Cre-responsive *Rosa-*DTA line driven by a Sertoli cell-specific *Amh-*cre. In addition, we also found impaired testicular vasculature in Sertoli cell-depleted fetal testes. A previous study reported that ablating Sertoli cells results in a reduction in total intratesticular vascular volume and the number of vascular branches (61). The treatment of PECAM1 monoclonal antibody blocks monocyte transmigration by 70–90% in some models (62). Thus, it is possible that the decrease of testicular monocyte number in Sertoli cell-depleted fetal testes may be due to the impaired testicular vasculature. Our previous study of macrophage-depleted fetal gonads by using *Cx3cr1*^cre^; *Rosa*-DTA mice revealed fewer endothelial cells migrating into the coelomic region of early fetal gonads. Therefore, whether Sertoli cells regulate testicular vasculature directly or via a secondary mechanism associated with monocytes needs to be further explored.

Testicular peritubular macrophages first appear in the testis upon the initiation of spermatogenesis at 2 weeks of age and are located in the myoid layer of the seminiferous tubule (23, 24), indicating the importance of cell niches composed of Sertoli cells and germ cells in regulating testicular macrophage differentiation and distribution. Our results revealed that the loss of Sertoli cell identity in *Dmrt1*^-/-^ postnatal testes, rather than germ cell loss, compromised the differentiation of peritubular macrophages, which subsequently had an abnormal interstitial localization. These defects may be due to chemokine and cytokine ligand-receptor systems that are required for macrophage differentiation, migration, and activation (63). Although monocyte chemoattractant protein-1 (MCP-1; official name CCL2) is detected in Sertoli cells (64) and its receptor, CCR2, is expressed in testicular peritubular macrophages (65), the number and localization of peritubular macrophages are completely normal in *Ccr2*^−/−^ mice (22, 24). This finding suggests that the MCP-1/CCR2 axis is unnecessary for peritubular macrophage differentiation and distribution in the testis. In addition, the CSF2 (also called GM-CSF)/CSF2R pathway induces M1 polarization of macrophages by increasing MHCII expression and antigen presentation capacity of macrophages (66). Testicular peritubular macrophages have a high level of MHCII expression and are classified as M1 macrophages (23, 26). However, whether the CSF2/CSF2R axis plays an important role in the regulation of Sertoli cells during peritubular macrophage development warrants further investigation. Therefore, more in-depth transcriptome analyses of Sertoli cells in *Dmrt1*^-/-^ testes may shed light on the underlying regulatory mechanisms of peritubular macrophage differentiation and localization.

Our previous study using a *Cx3cr1*-cre; *Rosa-*DTA-mediated approach showed that YS–derived macrophages regulate early fetal testis morphogenesis (21). In this study, we also found that E13.5 YS- macrophage-depleted gonads (using a CSF1R-antibody approach) have a large number of irregularly branched and fused testis cords. However, we observed numerous CD11b+ myeloid cells in the gonad-mesonephros border region. In our recently published results (25), gonads lacking *Mafb* and *Maf* (also called *c-Maf*), encoding transcription factors required for monocyte/macrophage development (67–69), also exhibited extensive supernumerary CD11b+ cells, and we demonstrated that these CD11b+ myeloid cells are monocytes; however, F4/80+ macrophages were not affected in *Maf* and *Mafb* mutant gonads.

Additionally, testis cords in *Maf* mutant gonads were smaller, but likely due to a significant decrease in the number of germ cells (25). However, our current results and our previous study (21) showed that macrophage-depleted gonads, either by using CSF1R antibody injection or a *Cx3cr1*^creER^; *Rosa*-DTA ablation method, have a normal number of germ cells. Thus, a disruption in the ratio between monocytes and macrophages may be a major cause of aberrant testis cords. Although testicular macrophages recovered or even exceeded their normal levels in CSF1R-injected E18.5 testes, as shown in previous reports on other fetal organs including liver, skin, kidney and lung (8), there was still a large number of CD11b+ monocytes with a decrease in the number of testis cords. Therefore, these results indicate that excess CD11b+ monocytes in fetal gonads may have a detrimental effect on testis cord morphogenesis and development, with the ratio of monocytes to macrophages being more important than macrophage quantity.

Macrophages and Leydig cells are in intimate contact throughout testis development and form intercytoplasmic digitations during puberty (70, 71), indicating essential roles of testicular macrophages in Leydig cell function. A body of evidence has shown that testicular macrophages promote testosterone production of Leydig cells in vitro (72); macrophage-deficient *Csf1^op/op^* mice are sub-fertile due to low testosterone levels (73, 74); and local depletion of testicular macrophages by intratesticularly injecting dichloromethylene diphosphonate-containing liposomes (Cl2MDP-lp) inhibits Leydig cell proliferation and regeneration (27, 75). Our results showed testicular macrophages stimulate Leydig cell steroidogenesis and proliferation in vitro via a co-culture model including varying ratios of Leydig cells and macrophages, which is more similar to the in vivo milieu of tight contact between Leydig cells and macrophages than previous experiments using conditioned culture media. Our previous findings showed that macrophage depletion by a diphtheria-toxin-mediated model reduced intratesticular testosterone levels (26). Therefore, testicular macrophages are a key regulator of Leydig cell steroidogenesis. After in vitro culture for a long period of time, despite the fact that testicular macrophages proliferated and reached substantial numbers, Leydig cells discontinued their high levels of testosterone secretion. Based on these findings regarding testosterone levels and macrophage proliferation, we propose that testosterone may inhibit testicular macrophage proliferation. Indeed, we found that androgen receptor (AR) was expressed in the membrane of testicular macrophages and had a nuclear localization in Leydig cells in vivo and in vitro (Fig. S10A and S10B), indicating that testosterone may inhibit testicular macrophage proliferation through non-genomic pathways. Testosterone also inhibits human monocyte/macrophage proliferation and induces the secretion of anti-inflammatory factors in vitro (76, 77). Adult testicular interstitial macrophages are in a non-proliferative state and highly express the anti-inflammatory factor IL-10 (23). Thus, non-proliferative testicular macrophages may stimulate testosterone synthesis via IL-10, but more research is needed. Our results also suggested that a significant increase in testicular macrophages suppressed testosterone production of Leydig cells during in vitro culture for long time. In addition, the reduction of steroidogenic gene expression in CSF1R-antibody-injected E18.5 testes may be related to the enhanced phagocytic activity of repopulated excess testicular macrophages and monocytes. In *Maf* mutant fetal gonads, we also found there were numerous monocytes with a downregulation of many steroidogenic genes (25). Therefore, a balance between Leydig cell steroidogenesis and testicular macrophage proliferation is critical for testis development and function.

## Materials and methods

### Mice

All mice used in this study were bred and housed under a 12-hour light/12-hour dark cycle and specific pathogen-free conditions in the Cincinnati Children’s Hospital Medical Center’s animal care facility, in compliance with institutional and National Institutes of Health guidelines. All experimental procedures were approved by the Institutional Animal Care and Use Committee (IACUC) of Cincinnati Children’s Hospital Medical Center (IACUC protocol # IACUC2021-0016). *Csf1r*-creER [Tg(Csf1r-Mer-iCre-Mer)^1Jwp^; JAX stock # 019098] (78), *Cx3cr1*^creER^ [*Cx3cr1*^tm2.1(creERT2)Jung^/J; JAX stock # 020940] (12), *Amh*-cre [Tg(Amh-cre)^8815Reb^/J; JAX stock # 033376] (46), *Rosa*-Tomato [Gt(ROSA)26Sor^tm14(CAG-tdTomato)Hze^/J; also called Ai14, JAX stock # 007914] (79), *Rosa*-DTA [Gt(ROSA)26Sor^tm1(DTA)Lky^/J; JAX stock # 009669] (80), *Cx3cr1*^GFP^ [*Cx3cr1*^tm1Litt^/J; JAX stock # 005582] (45), *Kit*^W^*/Kit*^W-v^ compound heterozygous males (WBB6F1/J- Kit^W^/Kit^W-v^/J; JAX stock #100410) and wild-type C57BL/6J (B6) mice (JAX stock #000664) were obtained from The Jackson Laboratory and maintained according to the instructions provided by The Jackson Laboratory. *Kit*^creER^ [*Kit*^tm2.1(cre/Esr1*)Jmol^/J] mice (81) (obtained from J. Molkentin, Cincinnati Children’s Hospital Medical Center, but are publicly available from Jackson Laboratories; JAX stock #032052), *Flt3*- cre mice (82) [Tg(Flt3-cre)#Ccb; obtained from K. Lavine, Washington University School of Medicine], and *Dmrt1*^-/-^ mice (49) [*Dmrt1*^tm1.1Zark^; obtained from D. Zarkower, University of Minnesota] were maintained on a B6 background. *Dnd1^Ter^* mice (53) were a gift from Blanche Capel (Duke University Medical Center); *Dnd1^Ter^* mice were originally on a 129T2/SvEmsJ background, but were backcrossed to C57BL/6J for 3 generations to eliminate the occurrence of testicular teratomas while maintaining complete germ cell depletion (83). *Dnd1^Ter/Ter^* homozygous mice generated by an intercross of *Dnd1*^Ter/+^ heterozygous mice were genotyped by using a Custom TaqMan SNP Genotyping Assay (Thermo Fisher #4332077). Embryonic time points were defined by timed matings, and noon on the day of the appearance of a vaginal plug was considered E0.5.

### Tamoxifen-induced creER activation

To label fetal-derived macrophages, *Cx3cr1*^creER^, *Kit*^creER^ and *Csf1r*-creER males were crossed with *Rosa*-Tomato reporter females; pregnant females were intraperitoneally injected at the embryonic stages indicated with 75 μg/g body weight of 4-hydroxytamoxifen (4OHT) (Sigma-Aldrich #H6278) and supplemented with 37.5 μg/g body weight of progesterone (Sigma-Aldrich #P0130) to prevent fetal abortions due to potential estrogenic effects of tamoxifen. For CreER induction in postnatal *Kit*^creER^*; Rosa*- Tomato animals, P4 and P5 mice were intraperitoneally injected with 50ug TAM (Sigma-Aldrich #T5648); adult mice were treated with 100 μg/g body weight of TAM on three consecutive days.

### Depletion of YS macrophages

To deplete yolk-sac-derived macrophages, pregnant B6 females were treated at E6.5 with a single intraperitoneal injection of 3 mg of anti-CSF1R mAb (Bio X Cell, clone AFS98) or rat IgG2a isotype control (Bio X Cell, clone 2A3) as previously described (8, 22).

### Isolation of primary testicular interstitial cells

Interstitial cells were isolated from adult (9-12 weeks old) wild-type B6 or heterozygous *Cx3cr1*^GFP^ mice following a previously described protocol (84). Briefly, decapsulated testes were digested with RPMI 1640 medium (Sigma #R5158) containing 0.25 mg/mL collagenase IV (Worthington #LS004186), 2% heat-inactivated fetal bovine serum (FBS; Thermo Fisher #16000044) in a 120-rpm shaker at 34°C for 10 min. FBS was added until the final concentration reached 10% to stop the enzymatic reaction. A single-cell suspension was obtained by removing seminiferous tubules with a 100 µm cell strainer (Corning #352360). The supernatant was pelleted, washed with RPMI 1640 medium, and incubated in ACK buffer (Life Technologies #A10492-01) for 3 minutes at room temperature to lyse erythrocytes.

### In vitro culture of testicular macrophages-enriched interstitial cells by MACS sorting

A dissociated single-cell suspension was passed through 30 µm nylon mesh (Pre-Separation Filters; Miltenyi Biotec #130-041-407) to remove cell clumps that could clog the column. After counting cell numbers, an appropriate number of cells was pipetted, which was defined as “pre-separation” interstitial cells. The remaining cells were pelleted and resuspended in 80 µl of MACS buffer (PBS pH 7.2, 0.5% BSA and 2 mM EDTA) and 20 µl anti-CD45 MicroBeads (Miltenyi Biotec #130-052-301) or 20 µl anti-F4/80 MicroBeads (Miltenyi Biotec #130-110-443) per 10^7^ total cells. The mixture was incubated in the dark at 4°C for 15 min, washed, and resuspended in MACS buffer. Cell suspension was applied onto the pre-balanced LS column (Miltenyi Biotec #130–097-679) that was placed in the magnetic Separator. Then the LS column was removed from the separator and the magnetically labeled cells were immediately flushed out, which were defined as the CD45+ or F4/80+ fraction. Unlabeled cells which passed through the LS column loading CD45 microbeads and LD column (Miltenyi Biotec #130-042-901) retaining weakly labeled cells were defined as the CD45- fraction. Each fraction was seeded in 24 well plates with or without glass coverslips and maintained in a humidified atmosphere (5% CO2, 95% air) at 37 °C for 3 days or 6 days in culture media (10% FBS in RPMI-1640).

### Flow cytometry

Each interstitial cell fraction in *in vitro* culture on day 3 or day 6 was digested with Accumax™ (STEMCELL Technologies #07921) for 5 min at 37 °C. Total cell number was determined in order to calculate the actual number of Leydig cells and macrophages based on percentages provided by the below flow cytometry analysis. Cell suspension was pelleted and washed with FACS buffer (2 mM EDTA, 2% FBS in PBS). Cells were incubated with an Fc blocker (anti CD32/16; BioLegend #101320) antibody for 10 min and then incubated with the respective mix of antibodies (Table S1) in FACS buffer for 30 min at 4°C. For viability staining, the labeled cells were washed and incubated with Zombie UV™ Fixable Viability Kit (BioLegend #423107) in FACS buffer before analysis. Flow cytometry analysis was performed on a BD Biosciences Fortessa flow cytometer. Viability staining and singlet profiling were always used to pre-gate cells. Data from two independent experiments were analyzed using FACS Diva software (BD Biosciences).

### Immunofluorescence

E12.5 and E13.5 gonads were dissected in PBS and fixed overnight in 4% paraformaldehyde (PFA) with 0.1% Triton X-100 at 4°C for whole-mount immunofluorescence as previously described (21). Testes of E14.5, E16.5, E18.5, P7, P18, P30, P60, P90 and P120 mice were dissected in PBS and fixed overnight in 4% PFA with 0.1% Triton X-100 at 4°C; after overnight fixation, testes were processed through a sucrose:PBS gradient (10%, 15%, 20% sucrose) and were embedded in OCT medium (Fisher Healthcare, #4585) at -80°C prior for cryosectioning, as previously reported (52). Then samples were washed several times in PBTx (PBS + 0.1% Triton X-100) and incubated in in blocking solution (PBTx + 10% FBS + 3% bovine serum albumin [BSA]) for 1-2 hours at room temperature. Primary antibodies used for immunofluorescence were diluted in blocking solution according to Table S1 and applied to samples overnight at 4°C. After several washes in PBTx, Alexa 488-, Alexa-555-, Alexa-647-, or Cy3-conjugated fluorescent secondary antibodies (Molecular Probes/Thermo Fisher or Jackson Immunoresearch, all at 1:500 dilution) and nuclear dye (2 mg/ml Hoechst 33342; Thermo Fisher #H1399) were diluted in blocking solution and applied to cryosections or interstitial cells for 1 hour and to whole gonads for 2-3 hours at room temperature. Samples were imaged either on a Nikon Eclipse TE2000 microscope (Nikon, Tokyo, Japan) with an OptiGrid structured illumination imaging system using Volocity software (PerkinElmer, Waltham, MA, USA) or on a Nikon A1 Inverted Confocal Microscope (Nikon, Tokyo, Japan).

Interstitial cell proliferation in *in vitro* culture was determined by the Click-it® EdU Alexa Fluor-488 Kit (Invitrogen/Thermo Fisher #C10337) according to the manufacturer’s instructions. For co-staining of EdU with anti-CYP11A1 antibody (85) for Leydig cells and anti-CD206 antibody for interstitial macrophages, interstitial cells plated on glass coverslips were incubated in culture media containing 10 µM EdU for 2 hours at 37°C and then fixed in 4% PFA for 10 min on ice followed by permeabilization with 0.5% Trion X- 100 for 20 min at room temperature. After several washes in in PBS containing 3% BSA, interstitial cells were incubated with Click-iT® reaction cocktail for 30 min at room temperature. The CYP11A1 and CD206 primary antibodies, as well as the corresponding secondary antibodies, were then incubated according to the cryosectioning protocol.

### Quantitative real-time PCR (qRT-PCR)

RNA extraction, cDNA synthesis, and qRT-PCR were performed as previously described (52). Briefly, total RNA was obtained with TRIzol Reagent (Invitrogen/Thermo Fisher #15596018). Genomic DNA was digested with DNase I (Amplification Grade; Thermo Fisher #18068015) to purify RNA. An iScript cDNA synthesis kit (BioRad #1708841) was used on 500ng of RNA for cDNA synthesis, as per manufacturer’s instructions. qRT-PCR was performed using the Fast SYBR Green Master Mix (Applied Biosystems/Thermo Fisher #4385616) on the StepOnePlus Real-Time PCR system (Applied Biosystems/Thermo Fisher). Expression levels were normalized to *Gapdh*. Relative fold changes in gene expression were calculated relative to controls using the 2^−△△Ct^ method. Primers used for qRT-PCR analysis are listed in Table S2.

### Hormone measurements

Testosterone measurements of cell culture media were performed by the University of Virginia Center for Research in Reproduction Ligand Assay and Analysis Core. Interstitial cell supernatants were collected on day 3 and day 6 of *in vitro* culture, centrifuged at 2,000 x g for 5 minutes at room temperature and stored at −80°C until analyzed.

### Cell counts and testis cord morphometric analyses

For all quantifications, images within a field of view 672 μm x 900 μm for each genotype or treatment were analyzed using ImageJ software (NIH). Testis cords of E13.5 and E18.5 XY gonads were visualized by staining with anti-AMH antibody. For E13.5 XY gonads, five testis cords of each whole-mount gonad (*n*=9 gonads from independent embryos from 3 independent control-antibody-injected litters; *n*=11 gonads from independent embryos from 4 independent anti-CSF1R-antibody-injected litters) were measured and averaged for height and width measurements. All testis cords of each E13.5 whole-mount gonad were counted and used for calculating the percentage of irregular testis cords. For E18.5 XY gonad, all testis cords in each image were counted from at least 3 different cryosections (cross-sections) per testis, from *n*=3 for control-antibody-injected or anti-CSF1R-antibody-injected testes, each from an independent biological replicate. For fetal or adult testicular macrophage and Tomato+ cell counting, the Cell Counter plug-in in ImageJ was used to manually count positive cells each image; quantifications were taken from at least 3 different cryosections per testis, from *n*=3 testes, each from an independent biological replicate. E14.5, E16.5 and E18.5 testis areas were outlined using the Polygon Selection tool, and the area in pixels was calculated with the Measure function; pixel area was then converted to square millimeters based on the pixel dimensions of each image.

### Statistical analyses

Statistical details of experiments, such as the exact value of *n*, what *n* represents, precision measures (mean ± SD), and statistical significance can be found in the Figure Legends. For qRT-PCR, statistical analyses were performed using Prism version 5.0 (GraphPad) and an unpaired, two-tailed Student t-test was performed to calculate *P* values based on ΔCt values. At least two gonads from independent embryos were pooled for each biological replicate (*<u>n</u>*≥3 biological replicates) in qRT-PCR analyses. For immunofluorescence, at least 3 sections from at least *n*=3 independent, individual animals (i.e., *n*=3 independent biological replicates) were examined for each time point (i.e., fetal stage or age) and/or experimental condition. For cell counting and morphometric analyses, sample sizes are listed above for each group. Graph results are shown as the value ± SD and statistical analyses were performed using an unpaired, two-tailed Student’s t-test.

## Data availability

The data that support the findings of this study are available from the authors on reasonable request.

## Acknowledgements

We thank Drs. Jeffery Molkentin, Kory Lavine, David Zarkower, and Blanche Capel for mice and Dr. Dagmar Wilhelm for anti-CYP11A1 antibody. This work was supported by Cincinnati Children’s Research Innovation and Pilot Funding and by National Institutes of Health (grants R35GM119458 and R01HD094698 to T.D.).

## Author contributions

X.G. conducted experiments, performed data analyses, and co-wrote and edited the manuscript. A.H. conducted experiments and edited the manuscript. T.D. supervised the project and co-wrote and edited the manuscript.

## References

1. Stefater JA, 3rd, Ren S, Lang RA, Duffield JS. Metchnikoff’s policemen: macrophages in development, homeostasis and regeneration. Trends Mol Med. 2011;17(12):743–752.

2. Wynn TA, Chawla A, Pollard JW. Macrophage biology in development, homeostasis and disease. Nature. 2013;496(7446):445–455.

3. Epelman S, Lavine KJ, Randolph GJ. Origin and functions of tissue macrophages. Immunity. 2014;41(1):21–35.

4. Ginhoux F, Guilliams M. Tissue-Resident Macrophage Ontogeny and Homeostasis. Immunity. 2016;44(3):439–449.

5. van Furth R, Cohn ZA, Hirsch JG, Humphrey JH, Spector WG, Langevoort HL. The mononuclear phagocyte system: a new classification of macrophages, monocytes, and their precursor cells. Bull World Health Organ. 1972;46(6):845–852.

6. Ginhoux F, Greter M, Leboeuf M, Nandi S, See P, Gokhan S, Mehler MF, Conway SJ, Ng LG, Stanley ER, Samokhvalov IM, Merad M. Fate mapping analysis reveals that adult microglia derive from primitive macrophages. Science. 2010;330(6005):841–845.

7. Gomez Perdiguero E, Klapproth K, Schulz C, Busch K, Azzoni E, Crozet L, Garner H, Trouillet C, de Bruijn MF, Geissmann F, Rodewald HR. Tissue-resident macrophages originate from yolk-sac-derived erythro-myeloid progenitors. Nature. 2015;518(7540):547–551.

8. Hoeffel G, Chen J, Lavin Y, Low D, Almeida FF, See P, Beaudin AE, Lum J, Low I, Forsberg EC, Poidinger M, Zolezzi F, Larbi A, Ng LG, Chan JK, Greter M, Becher B, Samokhvalov IM, Merad M, Ginhoux F. C-Myb(+) erythro-myeloid progenitor-derived fetal monocytes give rise to adult tissue-resident macrophages. Immunity. 2015;42(4):665–678.

9. Samokhvalov IM, Samokhvalova NI, Nishikawa S. Cell tracing shows the contribution of the yolk sac to adult haematopoiesis. Nature. 2007;446(7139):1056–1061.

10. Schulz C, Gomez Perdiguero E, Chorro L, Szabo-Rogers H, Cagnard N, Kierdorf K, Prinz M, Wu B, Jacobsen SE, Pollard JW, Frampton J, Liu KJ, Geissmann F. A lineage of myeloid cells independent of Myb and hematopoietic stem cells. Science. 2012;336(6077):86–90.

11. Sheng J, Ruedl C, Karjalainen K. Most Tissue-Resident Macrophages Except Microglia Are Derived from Fetal Hematopoietic Stem Cells. Immunity. 2015;43(2):382–393.

12. Yona S, Kim KW, Wolf Y, Mildner A, Varol D, Breker M, Strauss-Ayali D, Viukov S, Guilliams M, Misharin A, Hume DA, Perlman H, Malissen B, Zelzer E, Jung S. Fate mapping reveals origins and dynamics of monocytes and tissue macrophages under homeostasis. Immunity. 2013;38(1):79–91.

13. Orkin SH, Zon LI. Hematopoiesis: an evolving paradigm for stem cell biology. Cell. 2008;132(4):631–644.

14. Fijak M, Bhushan S, Meinhardt A. Immunoprivileged sites: the testis. Methods Mol Biol. 2011;677:459–470.

15. Fijak M, Meinhardt A. The testis in immune privilege. Immunol Rev. 2006;213:66–81.

16. Li N, Wang T, Han D. Structural, cellular and molecular aspects of immune privilege in the testis. Front Immunol. 2012;3:152.

17. Bhushan S, Tchatalbachev S, Lu Y, Frohlich S, Fijak M, Vijayan V, Chakraborty T, Meinhardt A. Differential activation of inflammatory pathways in testicular macrophages provides a rationale for their subdued inflammatory capacity. J Immunol. 2015;194(11):5455–5464.

18. Bhushan S, Theas MS, Guazzone VA, Jacobo P, Wang M, Fijak M, Meinhardt A, Lustig L. Immune Cell Subtypes and Their Function in the Testis. Front Immunol. 2020;11:583304.

19. Winnall WR, Muir JA, Hedger MP. Rat resident testicular macrophages have an alternatively activated phenotype and constitutively produce interleukin-10 in vitro. J Leukoc Biol. 2011;90(1):133–143.

20. Niemi M, Sharpe RM, Brown WR. Macrophages in the interstitial tissue of the rat testis. Cell Tissue Res. 1986;243(2):337–344.

21. DeFalco T, Bhattacharya I, Williams AV, Sams DM, Capel B. Yolk-sac-derived macrophages regulate fetal testis vascularization and morphogenesis. Proc Natl Acad Sci U S A. 2014;111(23):E2384–2393.

22. Lokka E, Lintukorpi L, Cisneros-Montalvo S, Makela JA, Tyystjarvi S, Ojasalo V, Gerke H, Toppari J, Rantakari P, Salmi M. Generation, localization and functions of macrophages during the development of testis. Nat Commun. 2020;11(1):4375.

23. Mossadegh-Keller N, Gentek R, Gimenez G, Bigot S, Mailfert S, Sieweke MH. Developmental origin and maintenance of distinct testicular macrophage populations. J Exp Med. 2017;214(10):2829–2841.

24. Wang M, Yang Y, Cansever D, Wang Y, Kantores C, Messiaen S, Moison D, Livera G, Chakarov S, Weinberger T, Stremmel C, Fijak M, Klein B, Pleuger C, Lian Z, Ma W, Liu Q, Klee K, Handler K, Ulas T, Schlitzer A, Schultze JL, Becher B, Greter M, Liu Z, Ginhoux F, Epelman S, Schulz C, Meinhardt A, Bhushan S. Two populations of self-maintaining monocyte-independent macrophages exist in adult epididymis and testis. Proc Natl Acad Sci U S A. 2021;118(1).

25. Li SY, Gu X, Heinrich A, Hurley EG, Capel B, DeFalco T. Loss of Mafb and Maf distorts myeloid cell ratios and disrupts fetal mouse testis vascularization and organogenesisdagger. Biol Reprod. 2021;105(4):958–975.

26. DeFalco T, Potter SJ, Williams AV, Waller B, Kan MJ, Capel B. Macrophages Contribute to the Spermatogonial Niche in the Adult Testis. Cell Rep. 2015;12(7):1107–1119.

27. Gaytan F, Bellido C, Aguilar E, van Rooijen N. Requirement for testicular macrophages in Leydig cell proliferation and differentiation during prepubertal development in rats. J Reprod Fertil. 1994;102(2):393–399.

28. Hales DB. Testicular macrophage modulation of Leydig cell steroidogenesis. J Reprod Immunol. 2002;57(1-2):3–18.

29. Hutson JC. Physiologic interactions between macrophages and Leydig cells. Exp Biol Med (Maywood). 2006;231(1):1–7.

30. Stremmel C, Schuchert R, Wagner F, Thaler R, Weinberger T, Pick R, Mass E, Ishikawa-Ankerhold HC, Margraf A, Hutter S, Vagnozzi R, Klapproth S, Frampton J, Yona S, Scheiermann C, Molkentin JD, Jeschke U, Moser M, Sperandio M, Massberg S, Geissmann F, Schulz C. Yolk sac macrophage progenitors traffic to the embryo during defined stages of development. Nat Commun. 2018;9(1):75.

31. Kierdorf K, Erny D, Goldmann T, Sander V, Schulz C, Perdiguero EG, Wieghofer P, Heinrich A, Riemke P, Holscher C, Muller DN, Luckow B, Brocker T, Debowski K, Fritz G, Opdenakker G, Diefenbach A, Biber K, Heikenwalder M, Geissmann F, Rosenbauer F, Prinz M. Microglia emerge from erythromyeloid precursors via Pu.1- and Irf8-dependent pathways. Nat Neurosci. 2013;16(3):273–280.

32. Plein A, Fantin A, Denti L, Pollard JW, Ruhrberg C. Erythro-myeloid progenitors contribute endothelial cells to blood vessels. Nature. 2018;562(7726):223–228.

33. Elsaid R, Meunier S, Burlen-Defranoux O, Soares-da-Silva F, Perchet T, Iturri L, Freyer L, Vieira P, Pereira P, Golub R, Bandeira A, Perdiguero EG, Cumano A. A wave of bipotent T/ILC-restricted progenitors shapes the embryonic thymus microenvironment in a time-dependent manner. Blood. 2021;137(8):1024–1036.

34. Theret M, Mounier R, Rossi F. The origins and non-canonical functions of macrophages in development and regeneration. Development. 2019;146(9).

35. Jing Y, Cao M, Zhang B, Long X, Wang X. cDC1 Dependent Accumulation of Memory T Cells Is Required for Chronic Autoimmune Inflammation in Murine Testis. Front Immunol. 2021;12:651860.

36. Pillay J, den Braber I, Vrisekoop N, Kwast LM, de Boer RJ, Borghans JA, Tesselaar K, Koenderman L. In vivo labeling with 2H2O reveals a human neutrophil lifespan of 5.4 days. Blood. 2010;116(4):625–627.

37. Grabert K, Sehgal A, Irvine KM, Wollscheid-Lengeling E, Ozdemir DD, Stables J, Luke GA, Ryan MD, Adamson A, Humphreys NE, Sandrock CJ, Rojo R, Verkasalo VA, Mueller W, Hohenstein P, Pettit AR, Pridans C, Hume DA. A Transgenic Line That Reports CSF1R Protein Expression Provides a Definitive Marker for the Mouse Mononuclear Phagocyte System. J Immunol. 2020;205(11):3154–3166.

38. Rae F, Woods K, Sasmono T, Campanale N, Taylor D, Ovchinnikov DA, Grimmond SM, Hume DA, Ricardo SD, Little MH. Characterisation and trophic functions of murine embryonic macrophages based upon the use of a Csf1r-EGFP transgene reporter. Dev Biol. 2007;308(1):232–246.

39. Ivanova NB, Dimos JT, Schaniel C, Hackney JA, Moore KA, Lemischka IR. A stem cell molecular signature. Science. 2002;298(5593):601–604.

40. Manova K, Nocka K, Besmer P, Bachvarova RF. Gonadal expression of c-kit encoded at the W locus of the mouse. Development. 1990;110(4):1057–1069.

41. Zhang M, Zhou H, Zheng C, Xiao J, Zuo E, Liu W, Xie D, Shi Y, Wu C, Wang H, Li D, Li J. The roles of testicular c-kit positive cells in de novo morphogenesis of testis. Sci Rep. 2014;4:5936.

42. Christensen JL, Weissman IL. Flk-2 is a marker in hematopoietic stem cell differentiation: a simple method to isolate long-term stem cells. Proc Natl Acad Sci U S A. 2001;98(25):14541–14546.

43. Buza-Vidas N, Woll P, Hultquist A, Duarte S, Lutteropp M, Bouriez-Jones T, Ferry H, Luc S, Jacobsen SE. FLT3 expression initiates in fully multipotent mouse hematopoietic progenitor cells. Blood. 2011;118(6):1544–1548.

44. Geissmann F, Jung S, Littman DR. Blood monocytes consist of two principal subsets with distinct migratory properties. Immunity. 2003;19(1):71–82.

45. Jung S, Aliberti J, Graemmel P, Sunshine MJ, Kreutzberg GW, Sher A, Littman DR. Analysis of fractalkine receptor CX(3)CR1 function by targeted deletion and green fluorescent protein reporter gene insertion. Mol Cell Biol. 2000;20(11):4106–4114.

46. Holdcraft RW, Braun RE. Androgen receptor function is required in Sertoli cells for the terminal differentiation of haploid spermatids. Development. 2004;131(2):459–467.

47. Yan RG, Li BY, Yang QE. Function and transcriptomic dynamics of Sertoli cells during prospermatogonia development in mouse testis. Reprod Biol. 2020;20(4):525–535.

48. Wang YQ, Cheng JM, Wen Q, Tang JX, Li J, Chen SR, Liu YX. An exploration of the role of Sertoli cells on fetal testis development using cell ablation strategy. Mol Reprod Dev. 2020;87(2):223–230.

49. Raymond CS, Murphy MW, O’Sullivan MG, Bardwell VJ, Zarkower D. Dmrt1, a gene related to worm and fly sexual regulators, is required for mammalian testis differentiation. Genes Dev. 2000;14(20):2587–2595.

50. Matson CK, Murphy MW, Sarver AL, Griswold MD, Bardwell VJ, Zarkower D. DMRT1 prevents female reprogramming in the postnatal mammalian testis. Nature. 2011;476(7358):101–104.

51. Huang S, Ye L, Chen H. Sex determination and maintenance: the role of DMRT1 and FOXL2. Asian J Androl. 2017;19(6):619–624.

52. Heinrich A, Potter SJ, Guo L, Ratner N, DeFalco T. Distinct Roles for Rac1 in Sertoli Cell Function during Testicular Development and Spermatogenesis. Cell Rep. 2020;31(2):107513.

53. Youngren KK, Coveney D, Peng X, Bhattacharya C, Schmidt LS, Nickerson ML, Lamb BT, Deng JM, Behringer RR, Capel B, Rubin EM, Nadeau JH, Matin A. The Ter mutation in the dead end gene causes germ cell loss and testicular germ cell tumours. Nature. 2005;435(7040):360–364.

54. Kurimoto K, Yabuta Y, Ohinata Y, Shigeta M, Yamanaka K, Saitou M. Complex genome-wide transcription dynamics orchestrated by Blimp1 for the specification of the germ cell lineage in mice. Genes Dev. 2008;22(12):1617–1635.

55. Squarzoni P, Oller G, Hoeffel G, Pont-Lezica L, Rostaing P, Low D, Bessis A, Ginhoux F, Garel S. Microglia modulate wiring of the embryonic forebrain. Cell Rep. 2014;8(5):1271–1279.

56. Wen Q, Zheng QS, Li XX, Hu ZY, Gao F, Cheng CY, Liu YX. Wt1 dictates the fate of fetal and adult Leydig cells during development in the mouse testis. Am J Physiol Endocrinol Metab. 2014;307(12):E1131–1143.

57. King KY, Goodell MA. Inflammatory modulation of HSCs: viewing the HSC as a foundation for the immune response. Nat Rev Immunol. 2011;11(10):685–692.

58. Lee YS, Kim MH, Yi HS, Kim SY, Kim HH, Kim JH, Yeon JE, Byun KS, Byun JS, Jeong WI. CX3CR1 differentiates F4/80(low) monocytes into pro-inflammatory F4/80(high) macrophages in the liver. Sci Rep. 2018;8(1):15076.

59. Kaur G, Thompson LA, Dufour JM. Sertoli cells--immunological sentinels of spermatogenesis. Semin Cell Dev Biol. 2014;30:36–44.

60. Kaur G, Vadala S, Dufour JM. An overview of a Sertoli cell transplantation model to study testis morphogenesis and the role of the Sertoli cells in immune privilege. Environ Epigenet. 2017;3(3):dvx012.

61. Rebourcet D, Wu J, Cruickshanks L, Smith SE, Milne L, Fernando A, Wallace RJ, Gray CD, Hadoke PW, Mitchell RT, O’Shaughnessy PJ, Smith LB. Sertoli Cells Modulate Testicular Vascular Network Development, Structure, and Function to Influence Circulating Testosterone Concentrations in Adult Male Mice. Endocrinology. 2016;157(6):2479–2488.

62. Gerhardt T, Ley K. Monocyte trafficking across the vessel wall. Cardiovasc Res. 2015;107(3):321–330.

63. Mantovani A, Sica A, Sozzani S, Allavena P, Vecchi A, Locati M. The chemokine system in diverse forms of macrophage activation and polarization. Trends Immunol. 2004;25(12):677–686.

64. Guazzone VA, Rival C, Denduchis B, Lustig L. Monocyte chemoattractant protein-1 (MCP- 1/CCL2) in experimental autoimmune orchitis. J Reprod Immunol. 2003;60(2):143–157.

65. Figueiredo AFA, Wnuk NT, Vieira CP, Goncalves MFF, Brener MRG, Diniz AB, Antunes MM, Castro-Oliveira HM, Menezes GB, Costa GMJ. Activation of C-C motif chemokine receptor 2 modulates testicular macrophages number, steroidogenesis, and spermatogenesis progression. Cell Tissue Res. 2021;386(1):173–190.

66. Cai H, Zhang Y, Wang J, Gu J. Defects in Macrophage Reprogramming in Cancer Therapy: The Negative Impact of PD-L1/PD-1. Front Immunol. 2021;12:690869.

67. Aziz A, Soucie E, Sarrazin S, Sieweke MH. MafB/c-Maf deficiency enables self-renewal of differentiated functional macrophages. Science. 2009;326(5954):867–871.

68. Moriguchi T, Hamada M, Morito N, Terunuma T, Hasegawa K, Zhang C, Yokomizo T, Esaki R, Kuroda E, Yoh K, Kudo T, Nagata M, Greaves DR, Engel JD, Yamamoto M, Takahashi S. MafB is essential for renal development and F4/80 expression in macrophages. Mol Cell Biol. 2006;26(15):5715–5727.

69. Nakamura M, Hamada M, Hasegawa K, Kusakabe M, Suzuki H, Greaves DR, Moriguchi T, Kudo T, Takahashi S. c-Maf is essential for the F4/80 expression in macrophages in vivo. Gene. 2009;445(1-2):66–72.

70. Hutson JC. Development of cytoplasmic digitations between Leydig cells and testicular macrophages of the rat. Cell Tissue Res. 1992;267(2):385–389.

71. Miller SC, Bowman BM, Rowland HG. Structure, cytochemistry, endocytic activity, and immunoglobulin (Fc) receptors of rat testicular interstitial-tissue macrophages. Am J Anat. 1983;168(1):1–13.

72. Yee JB, Hutson JC. Effects of testicular macrophage-conditioned medium on Leydig cells in culture. Endocrinology. 1985;116(6):2682–2684.

73. Cohen PE, Hardy MP, Pollard JW. Colony-stimulating factor-1 plays a major role in the development of reproductive function in male mice. Mol Endocrinol. 1997;11(11):1636–1650.

74. Cohen PE, Chisholm O, Arceci RJ, Stanley ER, Pollard JW. Absence of colony-stimulating factor-1 in osteopetrotic (csfmop/csfmop) mice results in male fertility defects. Biol Reprod. 1996;55(2):310–317.

75. Gaytan F, Bellido C, Morales C, Reymundo C, Aguilar E, Van Rooijen N. Effects of macrophage depletion at different times after treatment with ethylene dimethane sulfonate (EDS) on the regeneration of Leydig cells in the adult rat. J Androl. 1994;15(6):558–564.

76. Cutolo M, Capellino S, Montagna P, Ghiorzo P, Sulli A, Villaggio B. Sex hormone modulation of cell growth and apoptosis of the human monocytic/macrophage cell line. Arthritis Res Ther. 2005;7(5):R1124–1132.

77. Becerra-Diaz M, Song M, Heller N. Androgen and Androgen Receptors as Regulators of Monocyte and Macrophage Biology in the Healthy and Diseased Lung. Front Immunol. 2020;11:1698.

78. Qian BZ, Li J, Zhang H, Kitamura T, Zhang J, Campion LR, Kaiser EA, Snyder LA, Pollard JW. CCL2 recruits inflammatory monocytes to facilitate breast-tumour metastasis. Nature. 2011;475(7355):222–225.

79. Madisen L, Zwingman TA, Sunkin SM, Oh SW, Zariwala HA, Gu H, Ng LL, Palmiter RD, Hawrylycz MJ, Jones AR, Lein ES, Zeng H. A robust and high-throughput Cre reporting and characterization system for the whole mouse brain. Nat Neurosci. 2010;13(1):133–140.

80. Voehringer D, Liang HE, Locksley RM. Homeostasis and effector function of lymphopenia-induced “memory-like” T cells in constitutively T cell-depleted mice. J Immunol. 2008;180(7):4742–4753.

81. van Berlo JH, Kanisicak O, Maillet M, Vagnozzi RJ, Karch J, Lin SC, Middleton RC, Marban E, Molkentin JD. c-kit+ cells minimally contribute cardiomyocytes to the heart. Nature. 2014;509(7500):337–341.

82. Epelman S, Lavine KJ, Beaudin AE, Sojka DK, Carrero JA, Calderon B, Brija T, Gautier EL, Ivanov S, Satpathy AT, Schilling JD, Schwendener R, Sergin I, Razani B, Forsberg EC, Yokoyama WM, Unanue ER, Colonna M, Randolph GJ, Mann DL. Embryonic and adult-derived resident cardiac macrophages are maintained through distinct mechanisms at steady state and during inflammation. Immunity. 2014;40(1):91–104.

83. Cook MS, Munger SC, Nadeau JH, Capel B. Regulation of male germ cell cycle arrest and differentiation by DND1 is modulated by genetic background. Development. 2011;138(1):23–32.

84. Medar MLJ, Marinkovic DZ, Kojic Z, Becin AP, Starovlah IM, Kravic-Stevovic T, Andric SA, Kostic TS. Dependence of Leydig Cell’s Mitochondrial Physiology on Luteinizing Hormone Signaling. Life (Basel). 2020;11(1).

85. Svingen T, Francois M, Wilhelm D, Koopman P. Three-dimensional imaging of Prox1-EGFP transgenic mouse gonads reveals divergent modes of lymphangiogenesis in the testis and ovary. PLoS One. 2012;7(12):e52620.

