## Supplementary Data for "Testicular macrophages are recruited during a narrow time window by fetal Sertoli cells to promote organ-specific developmental functions"

**Supplementary Figure Legends**

**Supplementary Figure 1. YS-derived EMPs contribute to brain microglia and liver Kupffer cells.** (A-L) Images of brain (A-F) and liver (G-L) at E18.5 (A,C,E,G,I,K) or P90 (B,D,F,H,J,L) from *Csf1r*-creER; *Rosa*-Tomato mice exposed to 4OHT at E8.5 (A,B,G,H), E10.5 (C,D,I,J), or E12.5 (E,F,K,L). In all supplementary figures in this study, prime figures (e.g., A' relative to A) are higher-magnification images of the boxed regions in the image to their left. Arrowheads and arrows in I' and K' denote Tomato-expressing F4/80+ macrophages and Tomato-expressing F4/80-negative cells, respectively. Thin scale bar, 100 μm; thick scale bar, 25 μm. (M and N) Graphs showing quantification (*n*=3) of percent Tomato-expressing F4/80+ macrophages in E18.5 or P90 brain (M) or liver (N) from *Csf1r*-creER; *Rosa*-Tomato mice induced with 4OHT at various embryonic stages. Data are shown as mean +/- SD. **P*<0.05; ***P*<0.01; ****P*<0.001 (two-tailed Student's *t*-test).

**Supplementary Figure 2. Fetal *Csf1r*+ definitive progenitors give rise to myeloid cells.** (A-J) Images of E18.5 liver (A,B,E-H) and testis (C,D,I,J) from *Csf1r*-creER; *Rosa*-Tomato mice exposed to 4OHT at E10.5 (A,C,E,G,I) or E12.5 (B,D,F,H,J) stained for various hematopoietic markers such as CD45 (pan-immune marker), KIT (HSC marker), and GR1 (neutrophil/monocyte marker). Arrows in E' and F' denote Tomato+ KIT-expressing HSCs. Thin scale bar, 100 μm; thick scale bar, 25 μm.

**Supplementary Figure 3. Fetal *Csf1r*+ definitive progenitors give rise to lymphoid cells.** (A-I) Images of P30 (A,D,G) and P90 (B,C,E,F,H,I) spleen from *Csf1r*-creER; *Rosa*-Tomato mice exposed to 4OHT at E10.5 (A,B,D,E,G,H) or E12.5 (C,F,I) stained for various hematopoietic markers such as B220 (B cell marker), CD4 (T cell marker), and GR1 (neutrophil/monocyte marker). Thin scale bar, 100 μm; thick scale bar, 25 μm.

**Supplementary Figure 4. Fetal AGM-derived HSCs give rise to multiple immune cell populations.** (A-I) Images of E18.5 brain (A,B), E18.5 testis (C), P30 liver (D,F,H), and P30 spleen (E,G,I) from *Kit*$^{\mathrm{creER}}$; *Rosa*-Tomato mice exposed to 4OHT at E8.5 (A) or E10.5 (B-I) stained for various immune cell populations. Arrowhead in C' denotes Tomato-expressing IBA1+ macrophage and arrow in C' denotes Tomato-expressing

CD45+ IBA1-negative cell. Thin scale bar, 100 μm; thick scale bar, 25 μm.

**Supplementary Figure 5. Bone-marrow-derived HSCs do not contribute to testicular macrophages.** (A-L) Images of testes from P7 (A,D,G), P30 (B,E,H), P60 (C,F,I), and P120 (J-L) *Kit*$^{creER}$; *Rosa*-Tomato mice exposed to 4OHT at P4 and P5 (A-I) or P60 (J-L) stained for markers of various gonadal cell populations such as F4/80 (macrophages), CD45 (pan-immune cell marker), KIT (HSCs, differentiating spermatogonia, and adult Leydig cells), CYP11A1 (Leydig cells), PECAM1 (endothelial cells), CSF1R (adult interstitial macrophages), MHCII (adult peritubular macrophages), and SF1 (adult Leydig cells). Black arrows indicate Tomato+ Leydig cells; black arrowheads indicate Tomato+ endothelial cells; and white arrows indicate Tomato+ KIT+ differentiating spermatogonia. Thin scale bar, 100 μm; thick scale bar, 25 μm.

**Supplementary Figure 6. *Flt3*-expressing fetal liver progenitors make a minor contribution to brain microglia and liver Kupffer cells.** (A-F) Images of E14.5 liver (A,B), E18.5 liver (C,E), E18.5 brain (D) and E18.5 testis (F) from *Flt3*-cre; *Rosa*-Tomato embryos. Black arrowheads denote Tomato-expressing KIT+ CD45-negative cells; white arrowheads denote Tomato-expressing CD45+ KIT-negative cells; and white arrows denote Tomato-expressing CD45+ KIT+ cells. Thin scale bar, 100 μm; thick scale bar, 25 μm. (G) Graphs showing quantification (N=3) of percent Tomato-expressing F4/80+ macrophages in E18.5 brain or liver from *Flt3*-cre; *Rosa*-Tomato embryos. Data are shown as mean +/- SD.

**Supplementary Figure 7. Loss of Sertoli cell identity in *Dmrt1* KO testes inhibits peritubular macrophage differentiation and localization.** (A-L) Images of P7 (A,B,G,H), P18 (C,D,I,J), and P60 (E,F,K,L) gonads from XY *Dmrt1*$^{+/-}$ heterozygous control (A,C,E,G,I,K) and *Dmrt1*$^{-/-}$ KO (B,D,F,H,J,L) mice. SOX9 labels testicular Sertoli cells and FOXL2 labels ovarian granulosa cells. Dashed lines indicate seminiferous tubule boundaries. Arrowheads indicate MHCII+ peritubular macrophages with proper localization to the seminiferous tubule periphery and flattened morphology; arrows indicate MHCII+ macrophages with improper interstitial localization and/or aberrant round morphology. Thin scale bar, 100 μm; thick scale bar, 25 μm.

**Supplementary Figure 8. YS-derived EMPs give rise to early gonadal monocytes and macrophages.** (A-H) Images of E12.5 fetal testes from *Csf1r*-creER; *Rosa*-Tomato (A-D) and *Kit*$^{creER}$; *Rosa*-Tomato (E-H) embryos exposed to 4OHT at E8.5 (A,C,E,G) or E10.5 (B,D,F,H) stained for various markers such as F4/80 (macrophages), CD11b (myeloid cells such as monocytes and granulocytes), CSF1R (macrophages), PECAM1 (germ cells and endothelial cells), and DDX4 (germ cells). Arrowheads indicate Tomato-expressing F4/80+ or CSF1R+ macrophages; black arrowheads indicate Tomato-expressing DDX4+ germ cells; and white arrows indicate Tomato-expressing CD11b+ cells (likely monocytes). Thin scale bar, 100 μm; thick scale bar, 25 μm.

**Supplementary Figure 9. Flow cytometric analyses reveal purity and cellular composition of cell populations used for in vitro culture assays.** (A-C) Flow cytometry plots for 4 different cell populations (pre-separation, CD45-depleted, CD45-enriched, and F4/80-enriched) isolated from *Cx3cr1*$^{GFP/+}$ adult testes at day 0 (A) or C57BL/6J adult testes at day 3 (B) and day 6 (C) of in vitro culture. Cells were analyzed for markers of macrophages (*Cx3cr1*-GFP), adult interstitial macrophages (CD206), and adult Leydig cells (CD106/VCAM1). SSC denotes side scatter.

**Supplementary Figure 10. Androgen receptor (AR) is expressed in the cell membrane of adult interstitial macrophages.** Images of P90 adult C57BL/6J testis (A) and pre-separation population of testicular cells isolated from adult C57BL/6J testis after 2 days of in vitro culture (B). Arrows indicate F4/80+ or CD206+ interstitial macrophages that express AR; in contrast to Leydig cells (VCAM1+ or CYP17A1+), which express AR predominantly in the nucleus, macrophage expression of AR is localized more broadly throughout the cell and in the cell membrane. Thin scale bar, 100 μm; thick scale bar, 25 μm.

**Table S1. Primary antibodies used for immunofluorescence and flow cytometry.**

| Primary Antibody | Dilution | Source/Reference |
|---|---|---|
| Rat anti-F4/80 | 1:2,000 | AbD Serotec #MCA497RT |
| Rabbit anti-IBA1 | 1:1,000 | Wako #019–19741 |
| Goat anti-FOXL2 | 1:250 | Novus #100-1277 |
| Rabbit anti-CSF1R | 1:500 | Santa Cruz #sc-692 |
| Rat anti-MHCII | 1:500 | eBioscience #14–5321-81 |
| Rat anti-CD4 | 1:400 | BioLegend #100505 |
| Rat anti-B220 | 1:400 | eBioscience #14–0452-81 |
| Rat anti-GR1 | 1:500 | AbD Serotec #MCA2387T |
| Rat anti-NR5A1 (SF1) | 1:250 | Cosmo Bio #KAL-KO610 |
| Goat anti-PECAM1 | 1:250 | R&D #AF3628 |
| Rat anti-CD45 | 1:300 | BioLegend #103101 |
| Goat anti-C-KIT/SCFR | 1:400 | R&D #AF1356 |
| Goat anti-AMH/MIS | 1:500 | Santa Cruz #sc-6886 |
| Rabbit anti-SOX9 | 1:3,000 | Millipore #AB5535 |
| Rat anti-CD206 | 1:1,000 | AbD Serotec #MCA2235T |
| Goat anti-VCAM1 | 1:2,000 | R&D #AF643 |
| Rat anti-CD11b | 1:250 | BD Pharmingen #557395 |
| Rabbit anti-DDX4 | 1:1,000 | Abcam #ab13840 |
| Goat anti-CYP17A1 | 1:500 | Santa Cruz #sc-46081 |
| Rabbit anti-CYP11A1 | 1:500 | A gift from D. Wilhelm (1) |
| Alexa Fluor® 647 anti-mouse CD106 Antibody (VCAM1) (flow cytometry) | 1:100 | Biolegend #105712 |
| PE anti-mouse CD206 (MMR) Antibody (flow cytometry) | 1:200 | Biolegend #141705 |

**Table S2. Sequences of primers used for qRT-PCR analyses.**

| Gene name | Sequence (5' to 3') |
|---|---|
| *Cdh5* forward | TCCTCTGCATCCTCACTATCACA |
| *Cdh5* reverse | GTAAGTGACCAACTGCTCGTGAAT |
| *Adgre1 (F4/80)* forward | CCCCAGTGTCCTTACAGAGTG |
| *Adgre1 (F4/80)* reverse | GTGCCCAGAGTGGATGTCT |
| *Csf1r* forward | TGTCATCGAGCCTAGTGGC |
| *Csf1r* reverse | CGGGAGATTCAGGGTCCAAG |
| *Cx3cr1* forward | GAGTATGACGATTCTGCTGAGG |
| *Cx3cr1* reverse | CAGACCGAACGTGAAGACGAG |
| *Itgam* forward | CCACACTAGCATCAAGGGCA |
| *Itgam* reverse | AAGGGACACACTGACACCTG |
| *Ccr2* forward | ACACCCTGTTTCGCTGTAGG |
| *Ccr2* reverse | TGGCCTGGTCTAAGTGCTTG |
| *Itgb2* forward | GTGTCCCAGGAATGCACCAA |
| *Itgb2* reverse | TATCATCGGCTGGACAACCC |
| *Ptprc* forward | GGAGGACACAGCACATTGGA |
| *Ptprc* reverse | CCCCTGAGCAGCAATCATCA |
| *Cyp11a1* forward | TGGCCCCATTTACAGGGAGAA |
| *Cyp11a1* reverse | GGCATCTGAACTCTTAAACAGGA |
| *Cyp17a1* forward | CAGAGAAGTGCTCGTGAAGAAG |
| *Cyp17a1* reverse | AGGAGCTACTACTATCCGCAAA |
| *Hsd3b1* forward | CAAGTGTGCCAGCCTTCATCT |
| *Hsd3b1* reverse | TTCATGATTCTGTTCCTCGTGG |
| *Kit* forward | CATGGCGTTCCTCGCCT |
| *Kit* reverse | GCCCGAAATCGCAAATCTTT |
| *Pou5f1* forward | GGAGGAAGCCGACAACAATGA |
| *Pou5f1* reverse | TCCACCTCACACGGTTCTCAA |
| *Ddx4* forward | TACTGTCAGACGCTCAACAGGA |
| *Ddx4* reverse | ATTCAACGTGTGCTTGCCCT |
| *Amh* forward | CCACACCTCTCTCCACTGGTA |
| *Amh* reverse | GGCACAAAGGTTCAGGGGG |
| *Sox9* forward | GCGGAGCTCAGCAAGACTCTG |
| *Sox9* reverse | ATCGGGGTGGTCTTTCTTGTG |
| *Gapdh* forward | AGGTCGGTGTGAACGGATTTG |
| *Gapdh* reverse | TGTAGACCATGTAGTTGAGGTCA |

#### References

1. Svingen T, Francois M, Wilhelm D, Koopman P. Three-dimensional imaging of Prox1-EGFP transgenic mouse gonads reveals divergent modes of lymphangiogenesis in the testis and ovary. *PLoS One*. 2012;7(12):e52620.

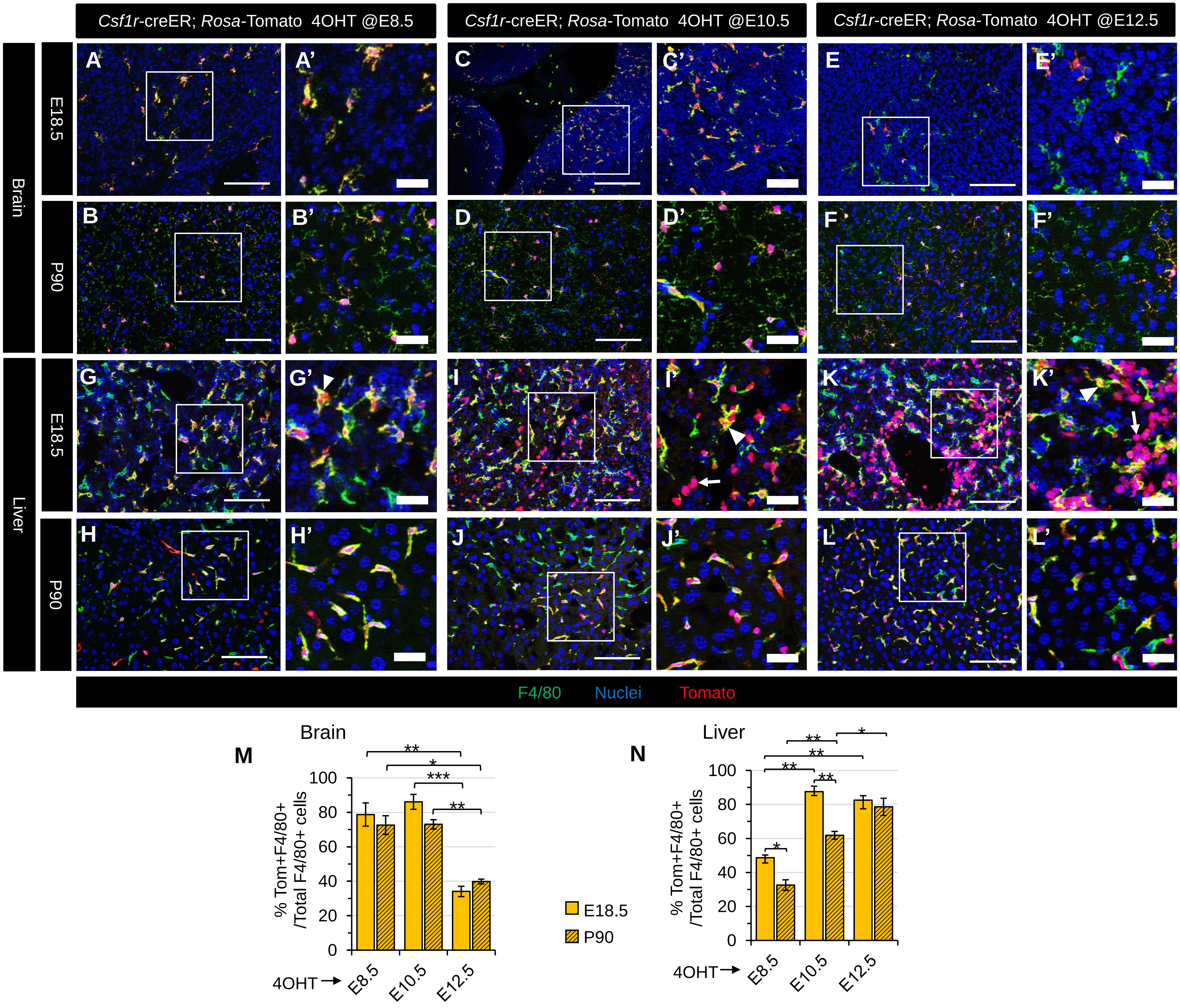


### Supplementary Figure S1

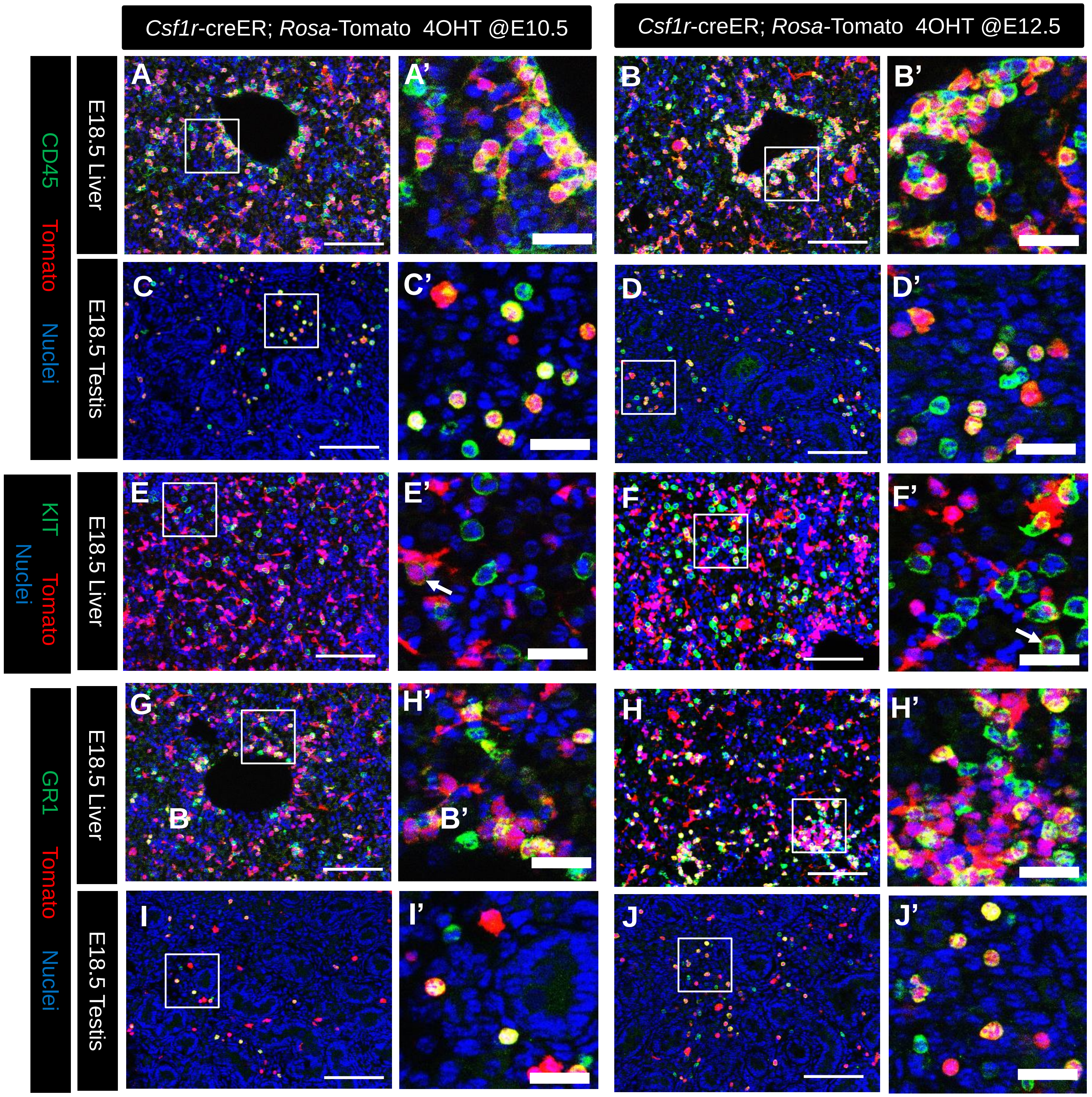


### Supplementary Figure S2

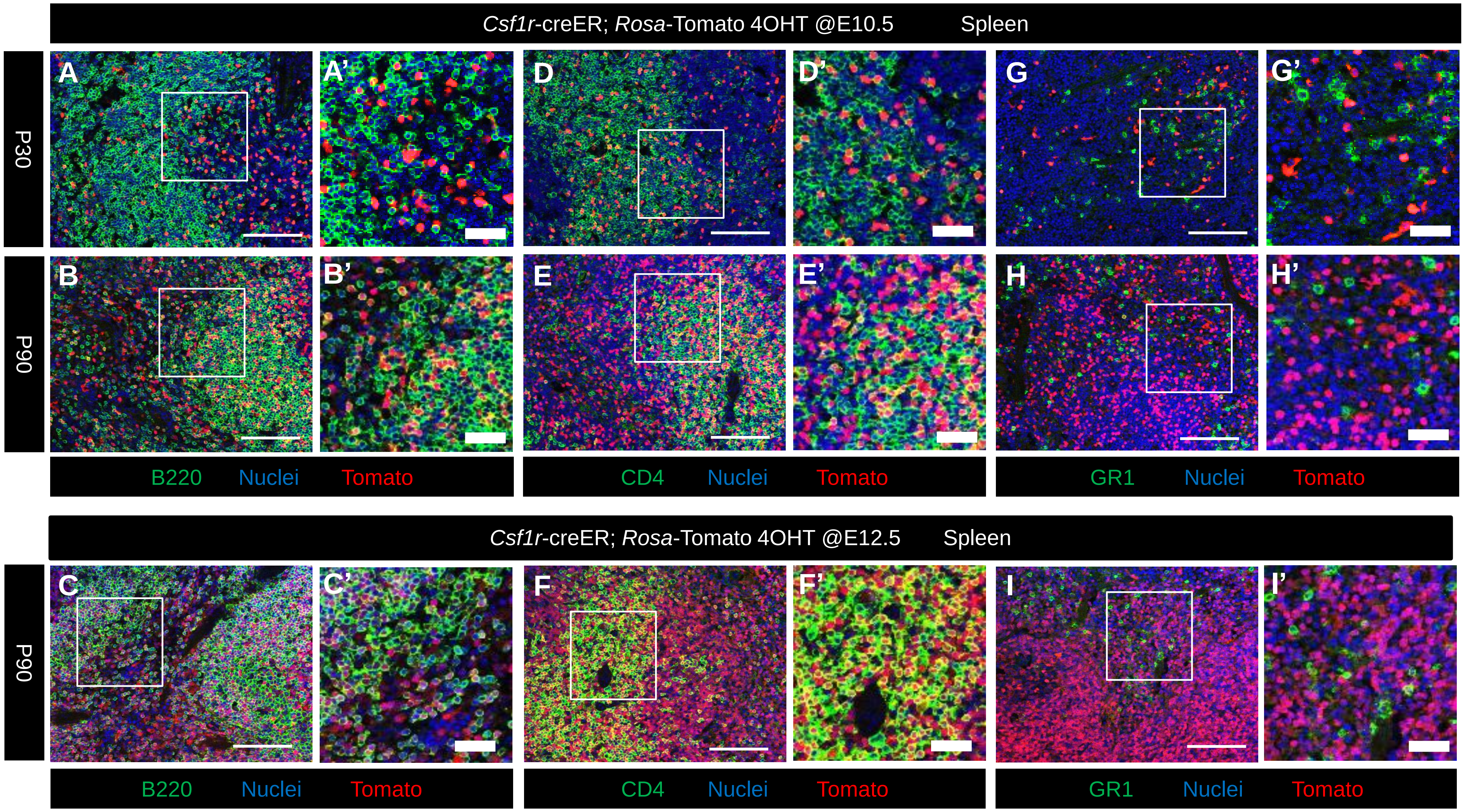


### Supplementary Figure S3

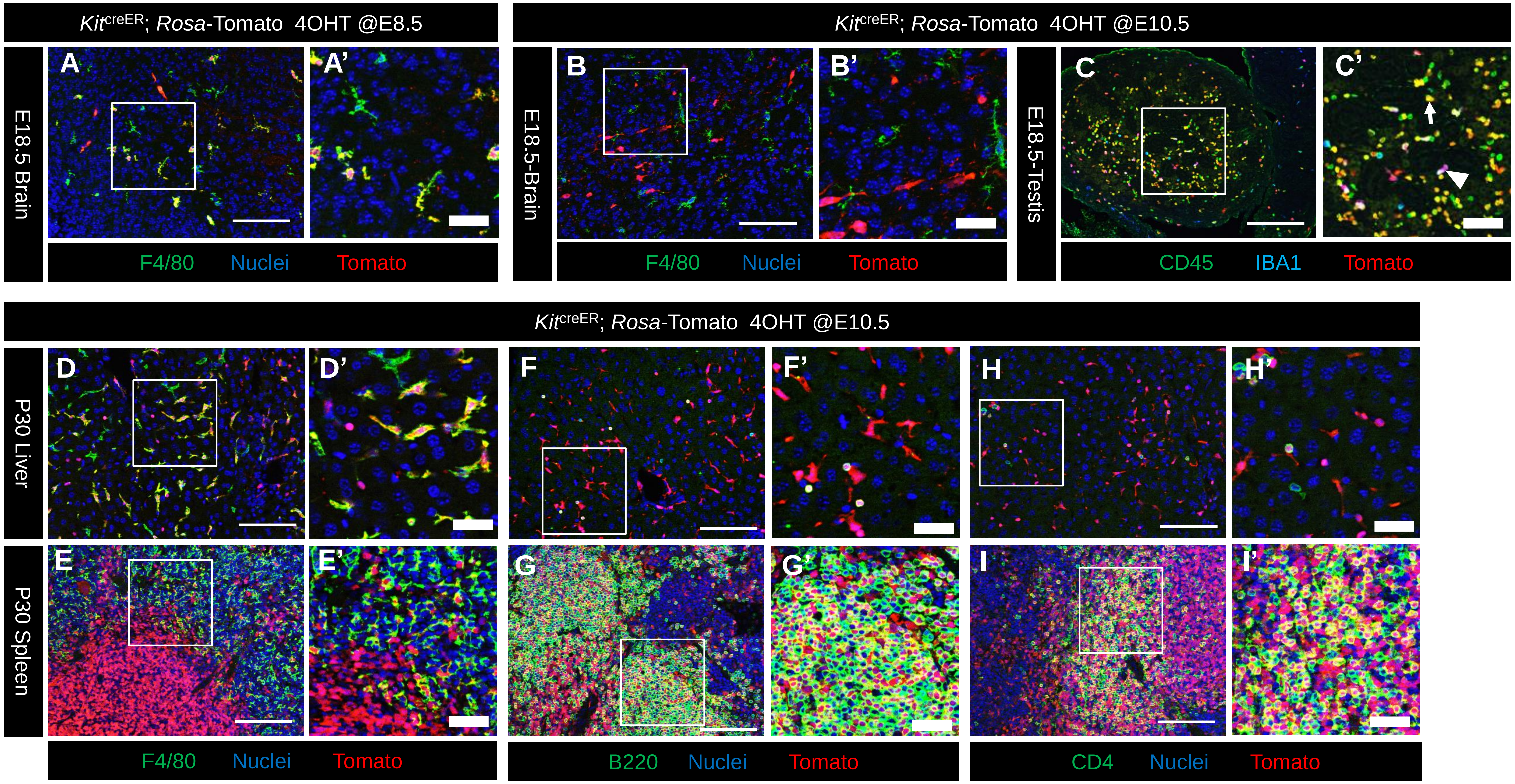


### Supplementary Figure S4

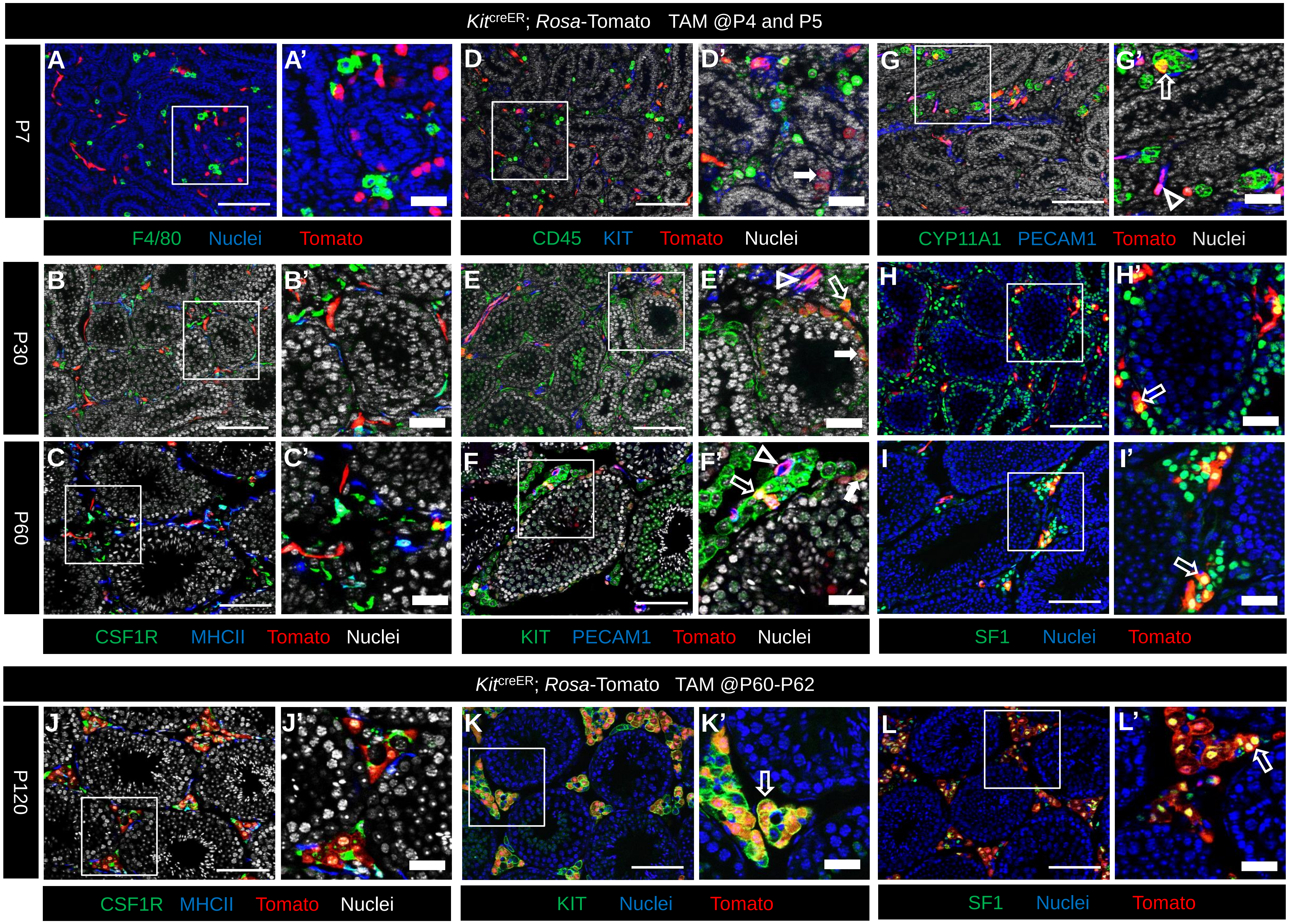


### Supplementary Figure S5

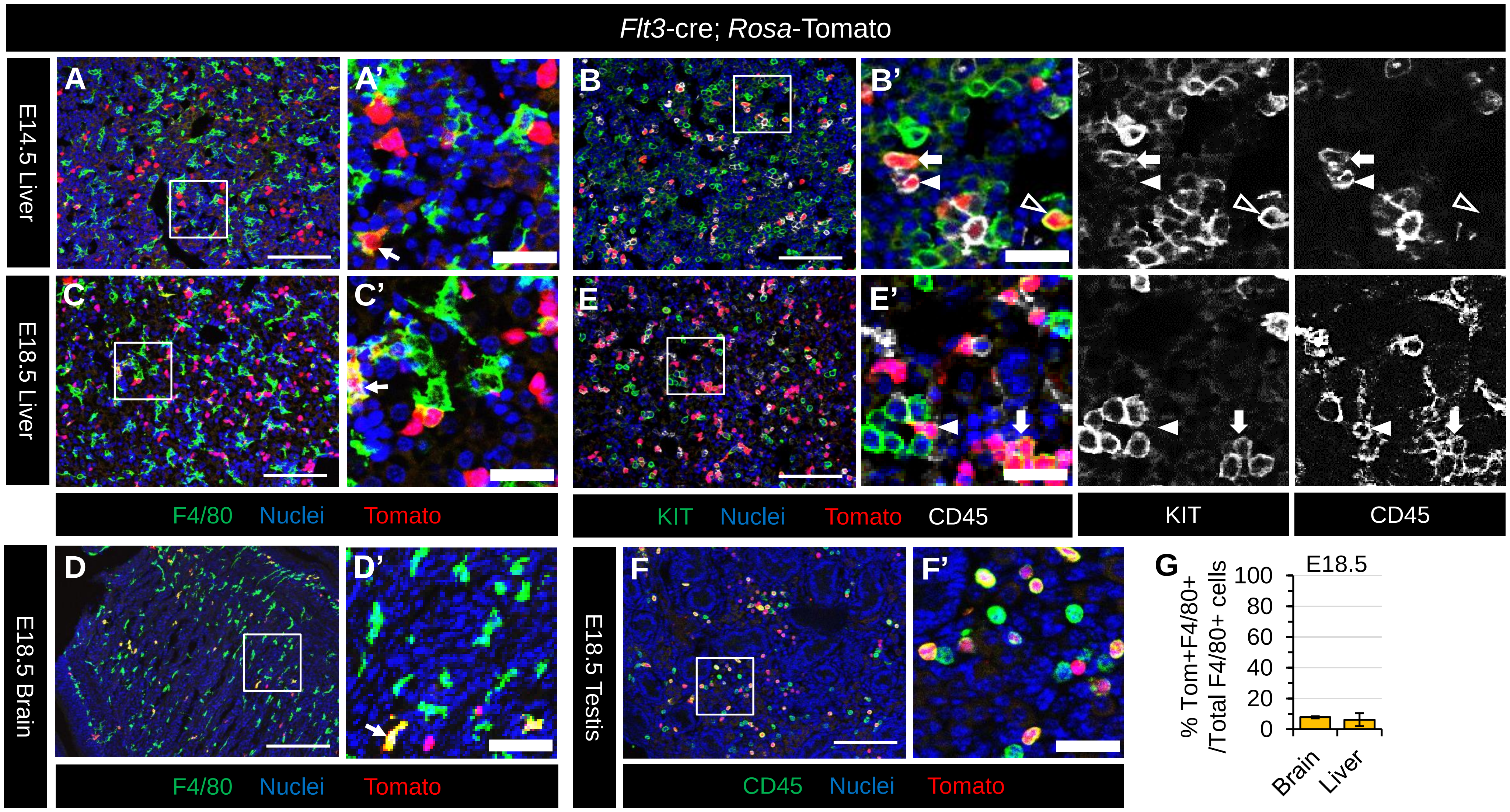


### Supplementary Figure S6

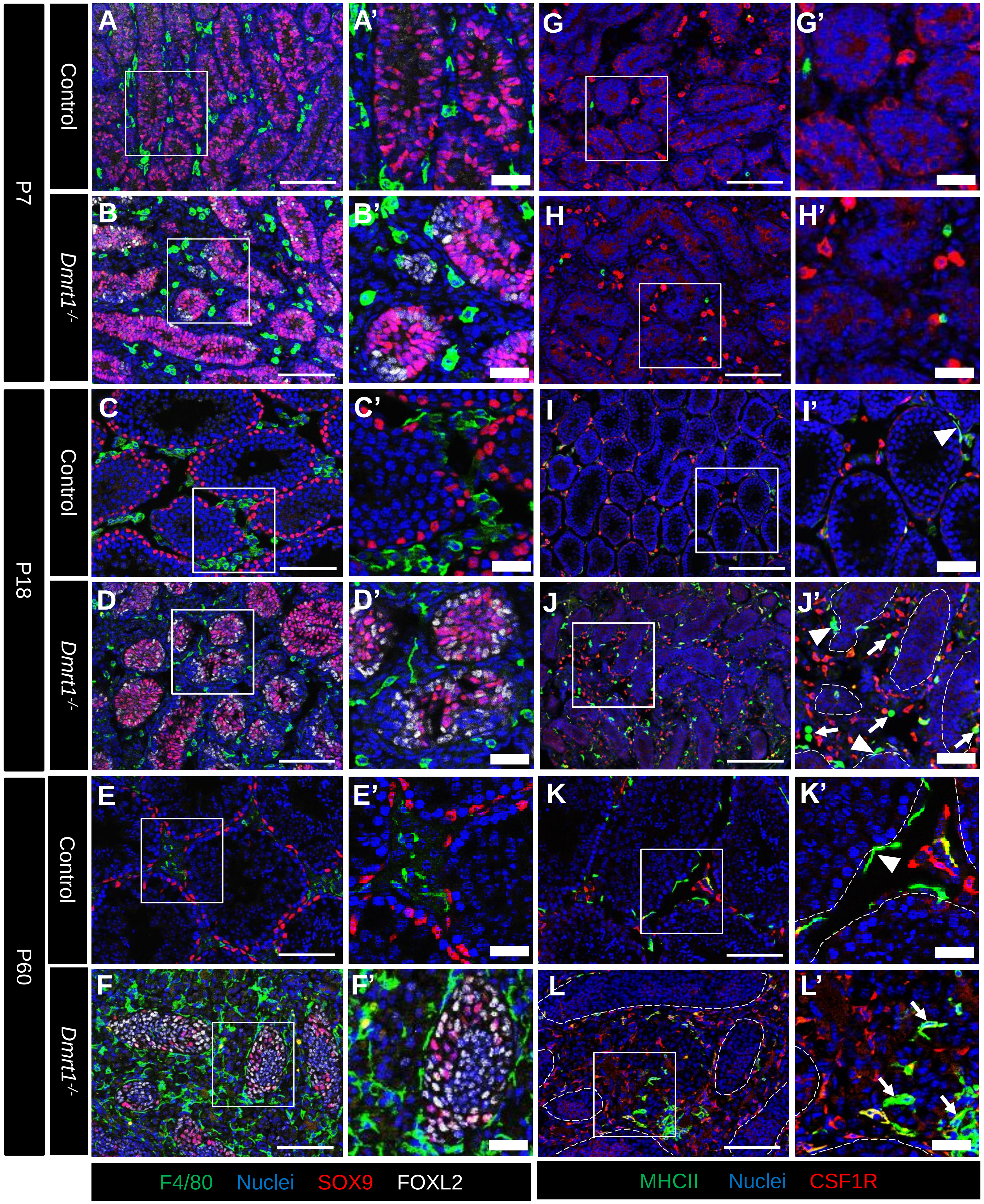


**Supplementary Figure S7**

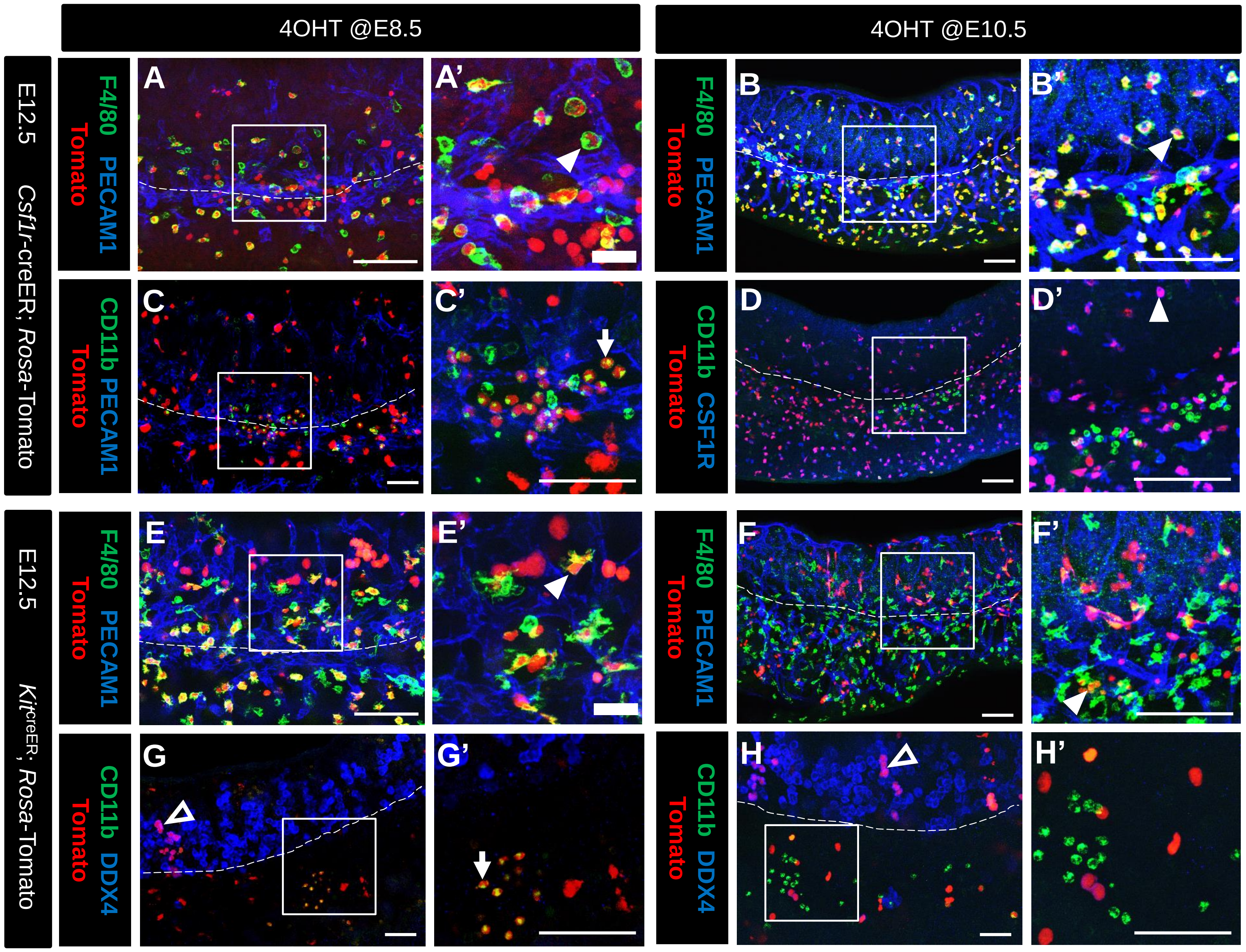


### Supplementary Figure S8

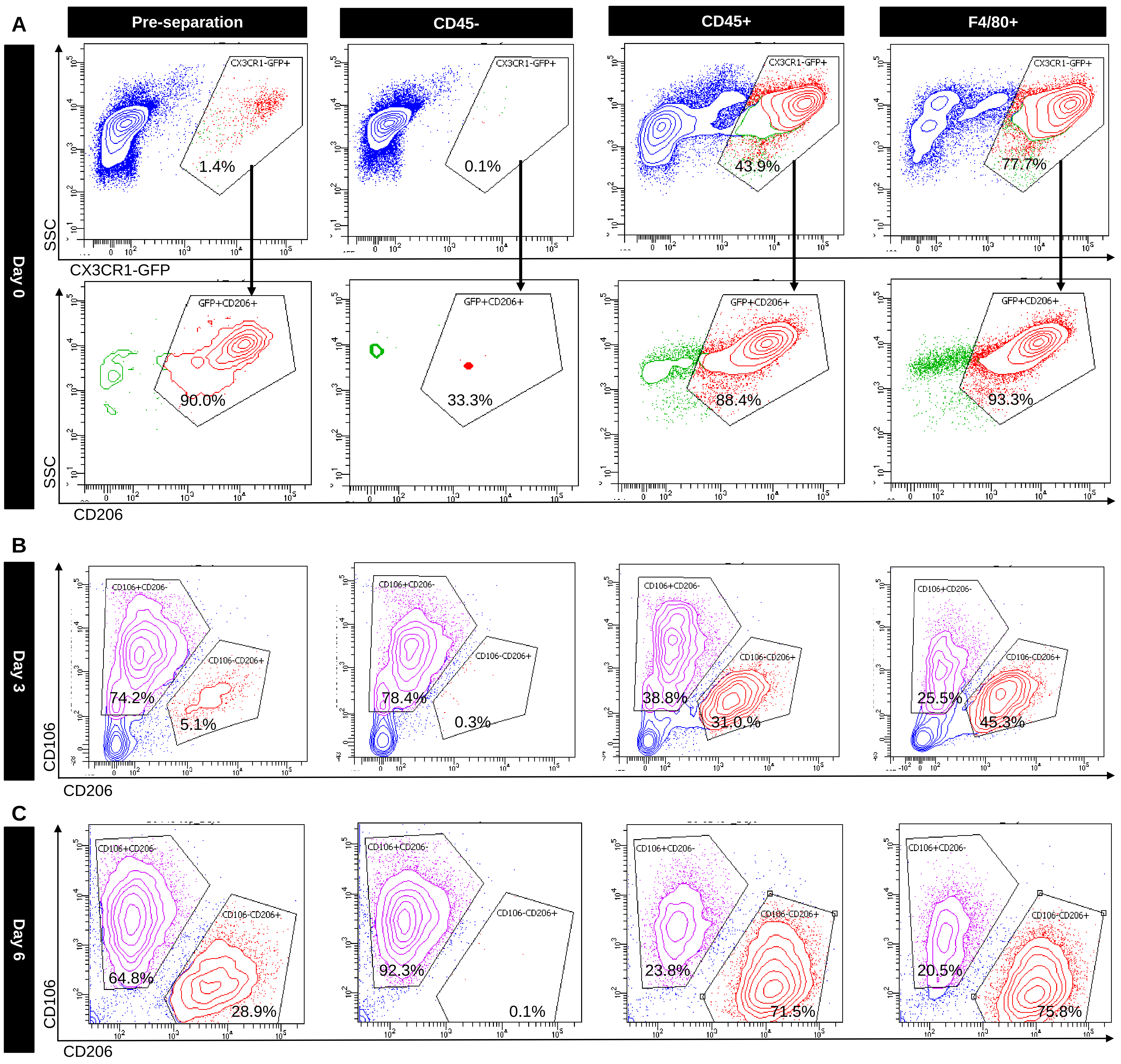


**Supplementary Figure S9**

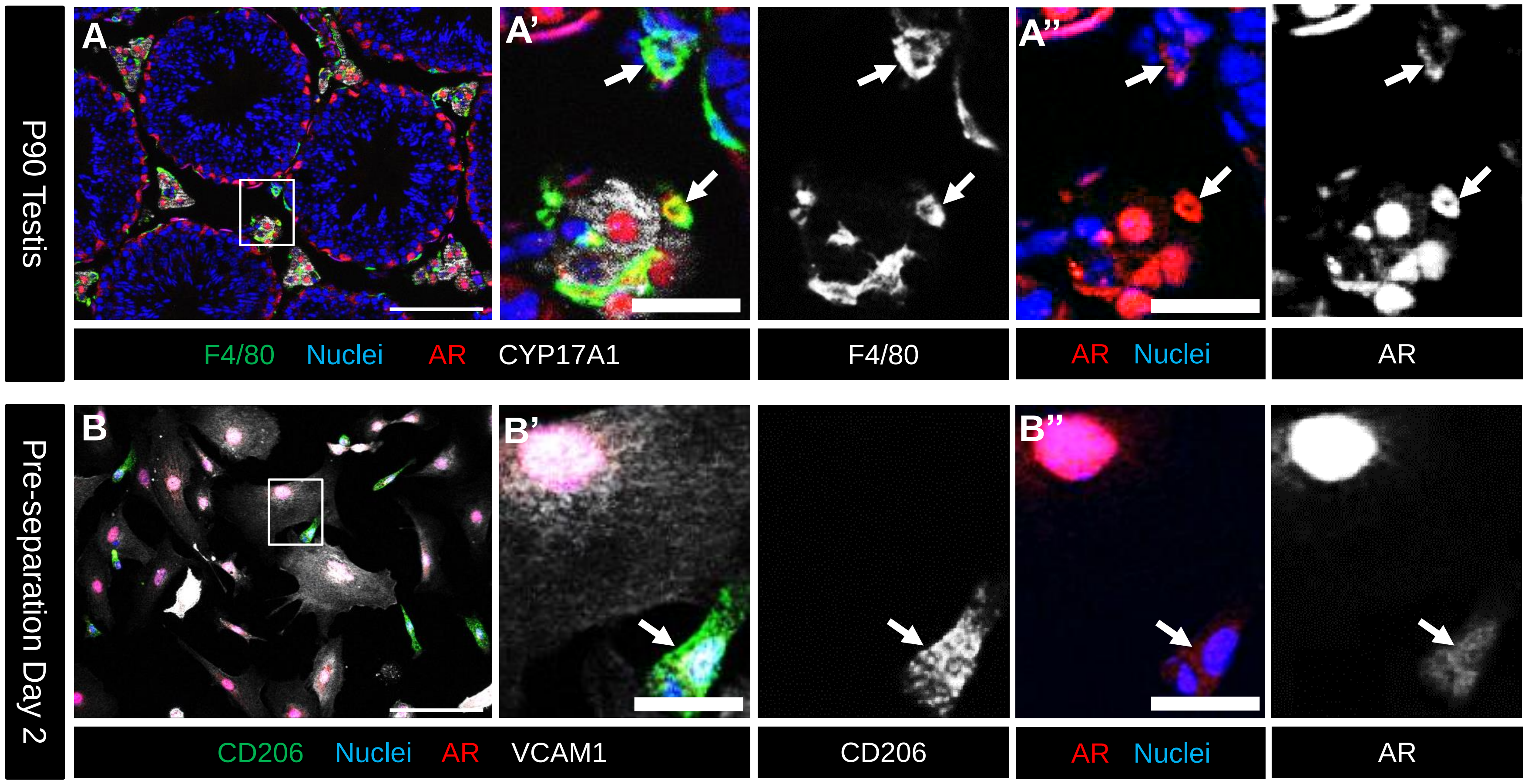


### Supplementary Figure S10
